# Allosteric activation of VCP, a AAA unfoldase, by small molecule mimicry

**DOI:** 10.1101/2023.10.02.560478

**Authors:** N.H. Jones, Q. Liu, L. Urnavicius, N.E. Dahan, L.E. Vostal, T.M. Kapoor

## Abstract

The loss of function of AAA (ATPases associated with diverse cellular activities) mechanoenzymes has been linked to diseases, and small molecules that activate these proteins can be powerful tools to probe mechanisms and test therapeutic hypotheses. Unlike chemical inhibitors that can bind a single conformational state to block enzyme activity, activator binding must be permissive to different conformational states needed for enzyme function. However, we do not know how AAA proteins can be activated by small molecules. Here, we focus on valosin-containing protein (VCP)/p97, a AAA unfoldase whose loss of function has been linked to protein aggregation-based disorders, to identify druggable sites for chemical activators. We identified VCP Activator 1 (VA1), a compound that dose-dependently stimulates VCP ATPase activity up to ∼3-fold. Our cryo-EM studies resulted in structures (∼2.9-3.5 Å-resolution) of VCP in apo and ADP-bound states, and revealed VA1 binding an allosteric pocket near the C-terminus in both states. Engineered mutations in the VA1 binding site confer resistance to VA1, and furthermore, modulate VCP activity to a similar level as VA1-mediated activation. The VA1 binding site can alternatively be occupied by a phenylalanine residue in the VCP C-terminal tail, a motif that is post-translationally modified and interacts with cofactors. Together, our findings uncover a druggable allosteric site and a mechanism of enzyme regulation that can be tuned through small molecule mimicry.

**Significance:** The loss of function of valosin-containing protein (VCP/p97), a mechanoenzyme from the AAA superfamily that hydrolyzes ATP and uses the released energy to extract or unfold substrate proteins, is linked to protein aggregation-based disorders. However, druggable allosteric sites to activate VCP, or any AAA mechanoenzyme, have not been identified. Here, we report cryo-EM structures of VCP in two states in complex with VA1, a compound we identified that dose-dependently stimulates VCP’s ATP hydrolysis activity. The VA1 binding site can also be occupied by a phenylalanine residue in the VCP C-terminal tail, suggesting that VA1 acts through mimicry of this interaction. Our study reveals a druggable allosteric site and a mechanism of enzyme regulation.

## Introduction

Proteins from the AAA superfamily of mechanoenzymes carry out essential functions across cell biology, including DNA replication, cytoskeleton remodeling, and membrane repair(1). AAA proteins are often hexameric, and use energy released during ATP hydrolysis in their AAA domains to carry out mechanical work on macromolecular substrates(2). The AAA mechanoenzyme VCP extracts or unfolds proteins associated with organelles or in complex cellular assemblies and its activity is central to numerous cellular processes, including endoplasmic reticulum-associated degradation (ERAD), autophagy, chromosome-associated degradation, and ribosome-associated quality control(3–5). Consistent with these important roles, VCP dysfunction is linked to several degenerative diseases, such as multisystem proteinopathy (MSP), amyotrophic lateral sclerosis (ALS), and vacuolar tauopathy, which shares characteristics with Alzheimer’s disease(6, 7). However, it is currently unclear which, if any, domain in VCP could be bound by a small molecule to stimulate its activity(8).

Structural data from crystallography and cryo-EM studies has provided key information about the domain organization of VCP and its overall mechanism(9–11). VCP assembles as a hexamer, and each VCP monomer consists of a globular N-terminal domain, two AAA domains (D1/D2), linker regions, and a disordered C-terminal tail (Fig. 1*A*)(12). Multiple lines of evidence indicate long-range allostery in VCP, from the N-domain through the D1 and D2 domains. For example, numerous protein cofactors regulate VCP function and direct it to specific, often ubiquitinated, substrates(13). Distinct cofactors can bind the N-terminal domain and/or the C-terminal tail, and can have inhibitory or stimulatory effects on VCP(14–16). Furthermore, small chemical changes such as point mutations and post-translational modifications across VCP’s domains can affect its activity(17–19). Finally, well-characterized chemical inhibitors of VCP have been reported and there is extensive structural data revealing how these compounds bind the enzyme D2 active site or an allosteric site (at the interface between the D1 and D2 domains)(8). However, the identification of putative allosteric binding sites for small molecule activators, primarily limited to computational methods, remains challenging(20).

**Fig. 1.**
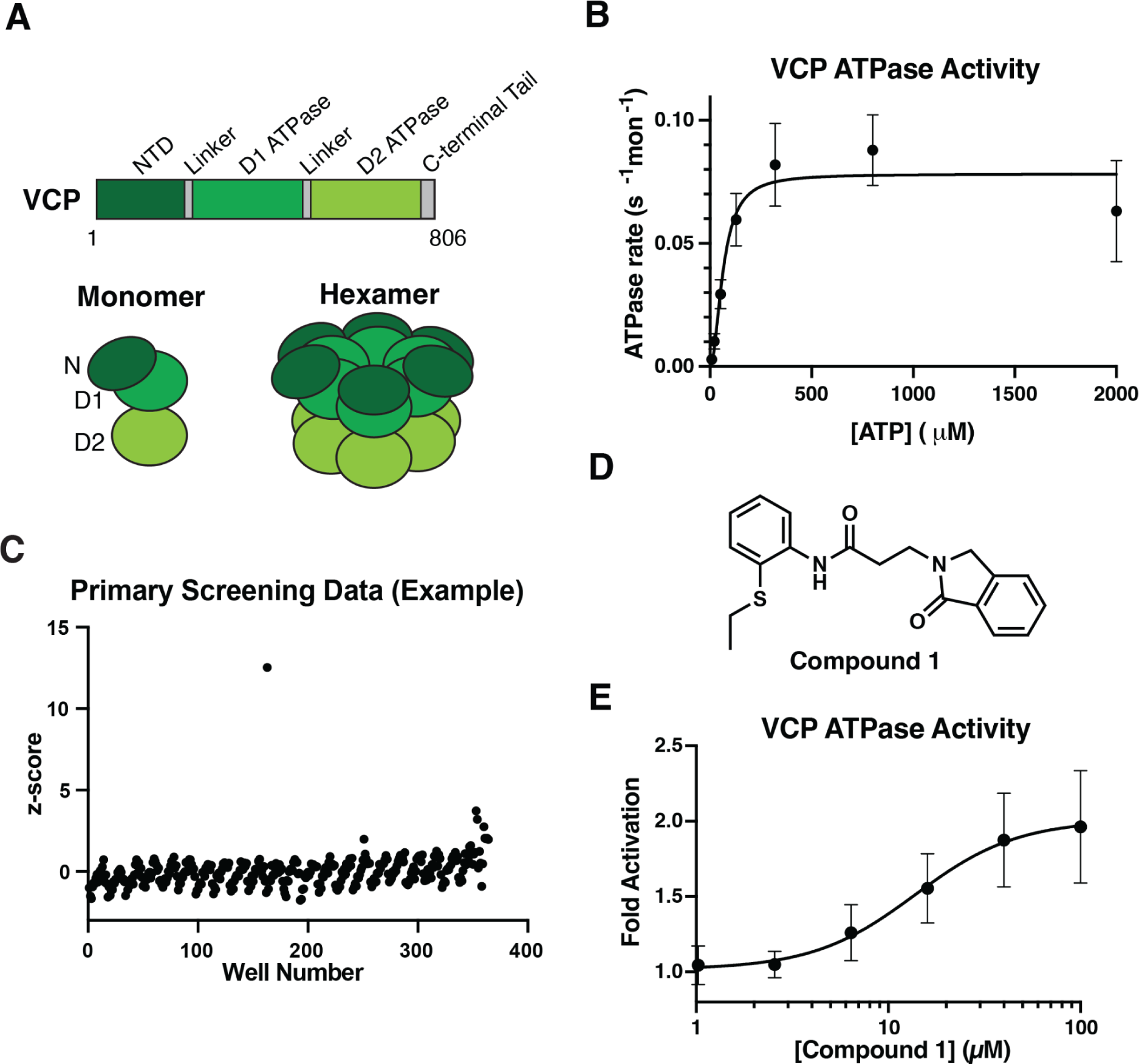
Screening for small molecule activators of VCP ATPase activity. (*A*) Schematics showing full-length VCP as a domain diagram (light gray box, not to scale, with the first and last residues and domains identified), a single monomer, and a hexamer. Colors indicate different domains. NTD = N-terminal domain. (*B*) ATP concentration dependence of the steady-state activity of VCP, analyzed using a Malachite Green assay. Rates were fit to the Michaelis-Menten equation for cooperative enzymes (mean +/− s.d., N = 6 independent experiments). (*C*) Calculated z-scores for compounds from a single screening plate tested in the ADPGlo assay. (*D*) Structure of Compound **1**. (*E*) Concentration-dependent activation of the ATPase activity of VCP by Compound **1** (100 µM ATP, 1 hr endpoint ADPGlo assay). Graph shows fold activation relative to DMSO control fit to a sigmoidal dose-response equation (mean +/− s.d., N = 9 independent experiments).

Activation of VCP may present a therapeutic opportunity, and small molecule activators of VCP have been recently reported(21–23). Currently, these compounds do not adhere to typical standards for physicochemical properties of drug-like molecules. For example, the activator SMER28 is a minimally substituted quinazoline, with a low molecular weight (∼260 Da), and may have multiple cellular targets(22). The binding site hypothesis for SMER28 is based on limited proteolysis-coupled mass spectrometry (LiP-MS) experiments and proposes interactions of this small molecule with the interface of the N-terminal/D1 domains. However, we lack the essential structural and mechanistic information needed to design drug-like activators of VCP.

In this manuscript, we discover a small molecule-binding allosteric regulatory site in VCP, a multi-domain homohexameric mechanoenzyme. We first identified a small molecule activator, which we named VCP Activator 1 (VA1), that dose-dependently stimulates the ATPase activity of VCP, saturating at ∼3-fold activation. Next, we used cryo-electron microscopy to characterize the structural basis of the VA1/VCP interaction, and found that VA1 binds in an allosteric pocket near the VCP D2 ATP-binding site in both the apo and ADP-bound states. Finally, we engineered point mutations in the VA1 binding pocket and show that they confer resistance to VA1 and stimulate or suppress VCP ATPase activity. Together, our data identify an important regulatory site that can be targeted by small molecule activators.

## Results

### Identification of a chemical activator of VCP ATPase activity

To identify a small molecule activator of VCP, we focused on the full-length protein (Fig. 1*A*). Recombinant VCP was purified using a published procedure with modifications(19), with 4 steps in total, which generated a recombinant untagged VCP construct at ∼93% purity (Fig. S1*A-B*) (see Materials and Methods). We characterized this construct using two assays. First, we carried out mass photometry experiments, which revealed a homogeneous species with a molecular weight of 533 +/− 6 kDa, consistent with a hexamer in solution (predicted hexamer molecular weight: 540 kDa) (Fig. S1*C*). Second, we measured kinetic parameters for ATP hydrolysis (k_cat_ = 0.084 +/− 0.006 s^-1^, K_1/2_ = 71 +/− 9 µM, mean +/− s.d., N = 6) (Fig. 1*B*) that are similar to previous reports(14, 17).

For the primary screen for VCP activators, we established an ATPase assay with optimized conditions that would result in ∼10-15% substrate conversion at ATP concentrations near the K_1/2_ (100 µM). Under these conditions (Z’: 0.85), screening ∼35,000 compounds (20 µM) resulted in a hit rate of 0.23% (cutoffs: z-score > 2, >1.2-fold activation) (Fig. 1*C*, example data from a single plate). Hit compounds were re-tested in three assays: 1) with the screening conditions, 2) against assay reagents in a counterscreen, and 3) at a range of doses. Validated dose-responsive hits represented five chemotypes. The most potent hit (“Compound **1**”) contains an isoindoline heterocycle and an aniline with a thioether substitution (Fig. 1*D*). Compound **1** was resynthesized and found to stimulate the ATPase activity of VCP in two different ATPase assays (maximum fold activation of 2.2 +/− 0.5, EC_50_ of 15 +/− 4 µM, mean +/− s.d., N = 9, 100 µM ATP, ADPGlo assay; similar results were obtained using a colorimetric assay) (Figs. 1*E* and S1*D*). At a higher ATP concentration (1 mM), dose-dependent activation of VCP by Compound **1** was also observed (maximum fold activation of 2.7 +/− 0.6, EC_50_ of 23 +/− 7 µM, mean +/− s.d., N = 3, 1 mM ATP, ADPGlo assay) (Fig. S1*E*). Finally, we tested different enzyme concentrations and did not observe a substantial effect of enzyme concentration on the EC_50_ of activation, consistent with a mechanism of activation that is not aggregation-mediated (Fig. S1*F*). Together, these data suggest that Compound **1** can stimulate the ATPase activity of VCP.

### Analyzing chemical activator binding to VCP

We next synthesized and tested analogs of Compound **1** (Compounds **2**-**6**, Table S1). Modification of the thioether substitution reduces VCP activation. Modifications to the linker region resulted in both improvements and losses in potency. For example, Compounds **4** and **5** represent an enantiopair, and the observed differences in their potencies suggest that VCP activation results from direct and specific interactions between the chemical scaffold and the enzyme. For additional studies, we focused on Compound **6** (hereafter, VCP Activator 1/VA1), which has more desirable properties (cLogP of 4.1 for Compound **5** versus 3.4 for VA1). VA1 shows a ∼3-fold increase in potency over Compound **1** (maximum fold activation of 3.0 +/− 0.3, EC_50_ of 4.1 +/− 1.1 µM, mean +/− s.d., N = 5, 100 µM ATP) (Figs. 2*A-B*).

**Fig. 2.**
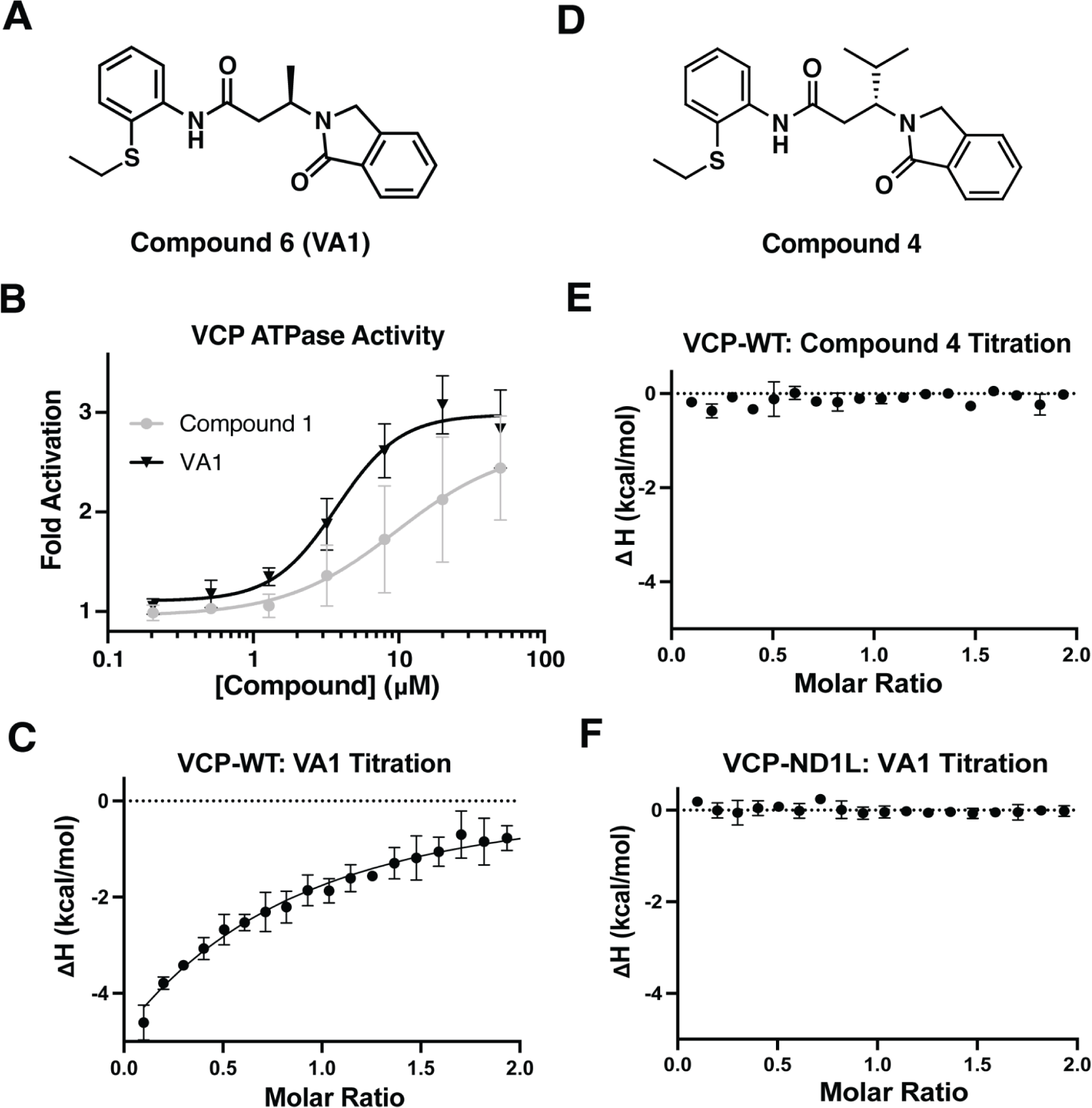
Analyzing direct binding of small molecule activators to VCP. (*A*) Structure of Compound **6** (VA1). (*B*) Concentration-dependent activation of the ATPase activity of VCP by Compound **1** and VA1 (100 µM ATP, 1 hr endpoint assay, ADPGlo). Graph shows fold activation relative to DMSO control fit to a sigmoidal dose-response equation (mean +/− s.d., N = 3 independent experiments for Compound **1**, N = 5 independent experiments for VA1). (*C*) Integrated data points and fitted binding curve used to determine K_d_ value from isothermal titration calorimetry (ITC)-based analysis of VCP in the presence of VA1 (mean +/− s.d., N = 3 independent experiments). (*D*) Structure of Compound **4**. (*E*) Integrated data points from ITC-based analysis of VCP in the presence of Compound **4** (mean +/− range, N = 2 independent experiments). (*F*) Integrated data points from ITC-based analysis of VCP-ND1L in the presence of VA1 (mean +/− range, N = 2 independent experiments).

To test whether VA1 makes direct interactions with VCP, we used isothermal titration calorimetry (ITC). The observed binding isotherm could be fit to a K_D_ of 27 +/− 4 µM (mean +/− s.d., N = 3) (Fig. 2*C*). Importantly, similar experiments with the inactive analog Compound **4** did not indicate any binding to VCP (Fig. 2*D-E*). We next examined if the N-terminal and D1 domains, which can bind protein co-factors that activate VCP(11, 14, 24–26), can be activated and bound by VA1. We purified a truncated construct with the VCP N-terminus and D1 domain (aa 1-472, hereafter VCP-ND1L) (Fig. S2*A*) (see Materials and Methods) and found its ATPase activity to be lower than that of the full-length protein, consistent with the role of the D2 domain as the primary source of hydrolysis activity (k_cat_ = 0.038 +/− 0.01 s^-1^, K_1/2_ = 48 +/− 8 µM, mean +/− range, N = 2) (Fig. S2*B*)(27). Interestingly, VA1 did not stimulate the ATPase activity of VCP-ND1L (up to 50 µM, ADPGlo assay) (Fig. S2*C*). Additionally, we did not observe binding of VA1 to VCP-ND1L in ITC experiments under similar conditions to those used for full-length VCP (Fig. 2*F*). Together, these data suggest that VA1 stimulates VCP ATPase activity through direct interactions.

### Structural analysis of VA1 binding VCP

To examine how VA1 binds VCP, we employed single-particle cryo-electron microscopy (cryo-EM). We incubated VCP with VA1 (200 µM), prepared grids, and observed homogenous, monodisperse particles in our micrographs (Fig. S2*D*; see Materials and Methods). Autopicked particles were cleaned by multiple rounds of 2D classification yielding class averages that resemble the VCP hexamer (Figs. S2*E* and S3*A*; see Materials and Methods). 3D classification to balance views and 3D refinement without symmetry of the cleaned particles from these classes resulted in an overall map of the VCP hexamer (Figs. 3*A* and S3*A*). The core resolved to ∼3.5-4.0 Å, revealing side chain detail, and the periphery resolved to ∼5.5-8 Å, revealing secondary structure details (Fig. 3*A*).

**Fig. 3.**
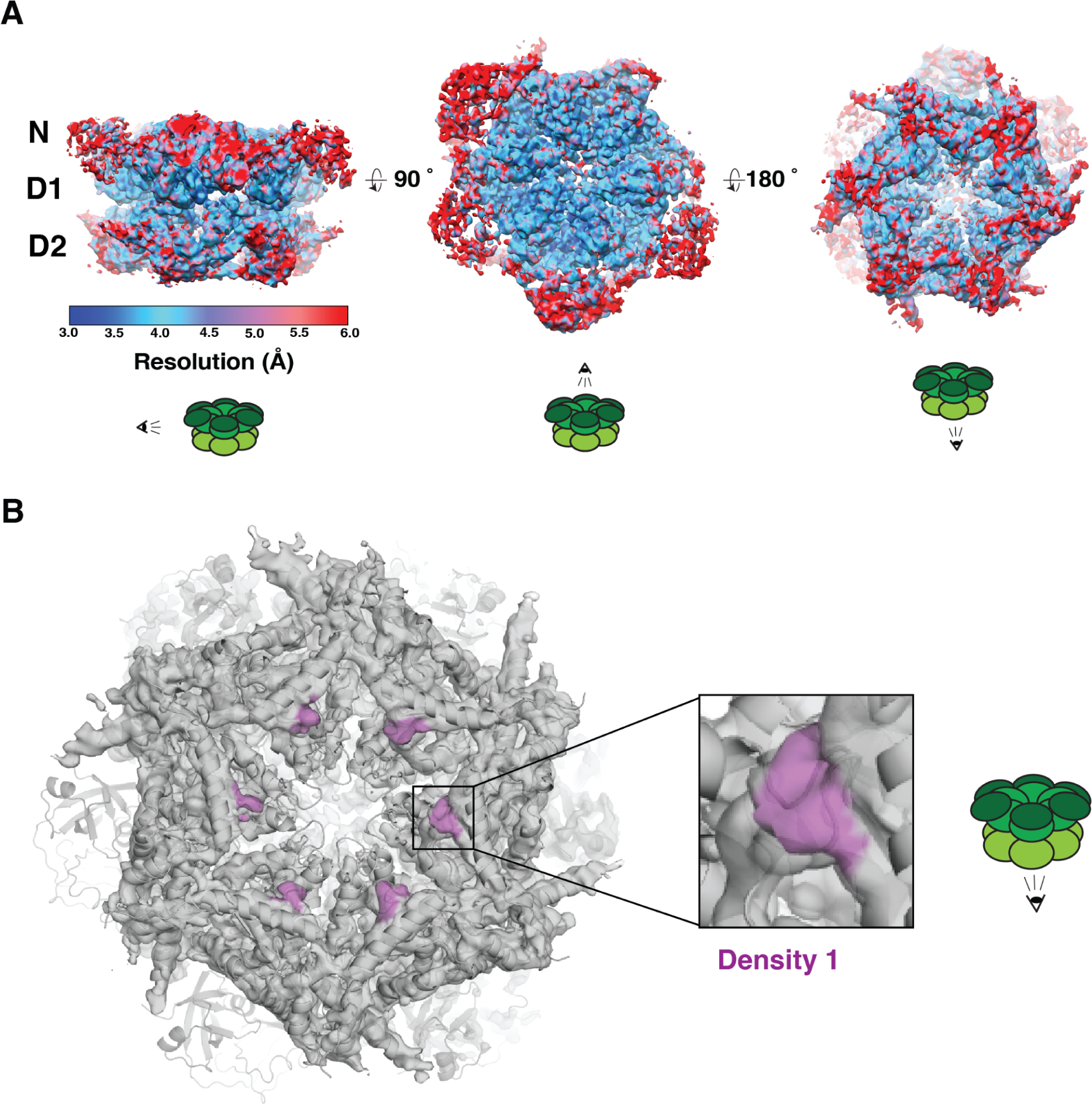
Cryo-EM analysis of VCP in the presence of VA1. (*A*) Three views of the overall VCP density analyzed by ResMap(53), showing a resolution distribution from 3.0 to 6.0 Å. (*B*) Molecular model of VCP (cartoon representation) in the density map (left, semi-transparent surface). Inset shows extra density (magenta) that could not be assigned to VCP.

We used rigid body docking to fit individual domains of a published ADP-bound VCP structure (PDB: 5FTL)(10), in addition to the C-terminal tail resolved in a peptide substrate-bound VCP structure (PDB: 7LN6)(11), into our map, with the D1 and D2 domains at the core, and the N-terminal domains and C-terminal tails, which are more dynamic(10, 11), at the periphery (Fig. 3*A-B*). Close inspection of our map revealed a density in the D2 domain in each monomer that could not be assigned to the VCP backbone or sidechains, or bound nucleotides (hereafter, “Density 1”) (Fig. 3*B*). Density 1 is localized between adjacent D2 nucleotide binding sites and the central pore, in a region that is well-resolved (Fig. 3*A-B*). Together, our findings with VCP-ND1L and our structural data suggest that Density 1 could represent VA1 (Fig. 3*B*).

### VA1 binds VCP in apo and ADP-bound states of the D2 domain

To better characterize the binding site containing Density 1, we carried out additional processing steps, focusing on the D2 domain, as signal subtraction from raw images followed by focused 3D refinement has proven useful for resolution improvements in other structures of large complexes(28–31). We additionally applied symmetry to obtain a map that resolved to ∼3.5 Å, and we could observe detailed interactions between Density 1 and nearby amino acid side chains. Only minor adjustments (real space sphere refinement within Coot; see Materials and Methods) to the docked VCP coordinates for residue side chains surrounding Density 1 were required (Fig. S4*A*).

To fit a model for VA1 in Density 1, we used computational modeling (Maestro Macromodel, Schrodinger LLC), which generated a bent conformation of the compound, placing the isoindoline and aniline proximal to each other. This shape of VA1 matches Density 1; the density is stronger for the aromatic rings than for the smaller amide linker, which is expected at this resolution (Figs. 4*A* and S4*A*). To address the weaker density for the amide linker, we focused on a small fraction (<5%) of particles corresponding to 2D classes that appeared to represent a VCP dodecamer in our cryo-EM data, a dimer of hexamers that has been observed in other cryo-EM studies of VCP (Figs. S2*E* and S3*A*)(32–35). Signal subtraction and focused refinement with D6 symmetry on the D2 domains resulted in a map resolved to ∼2.9 Å (Fig. S3*A*). Gratifyingly, we observed a density that agrees with Density 1 from the VCP hexamer map, and additionally shows continuous density between the isoindoline and aniline (Figs. 4*B* and S4*B*), matching our VA1 binding model. The narrower side of Density 1 accommodates the thioether substitution on the aniline and orients into a cleft formed by the Arg625/Asp627 loop and Tyr517 (positioned at the end of strand 1 of the β-sheet) (Fig. 4*C*). In this binding mode, the isoindoline is positioned near Met757 and Phe758 in the C-terminal helix (Fig. 4*D*).

**Fig. 4.**
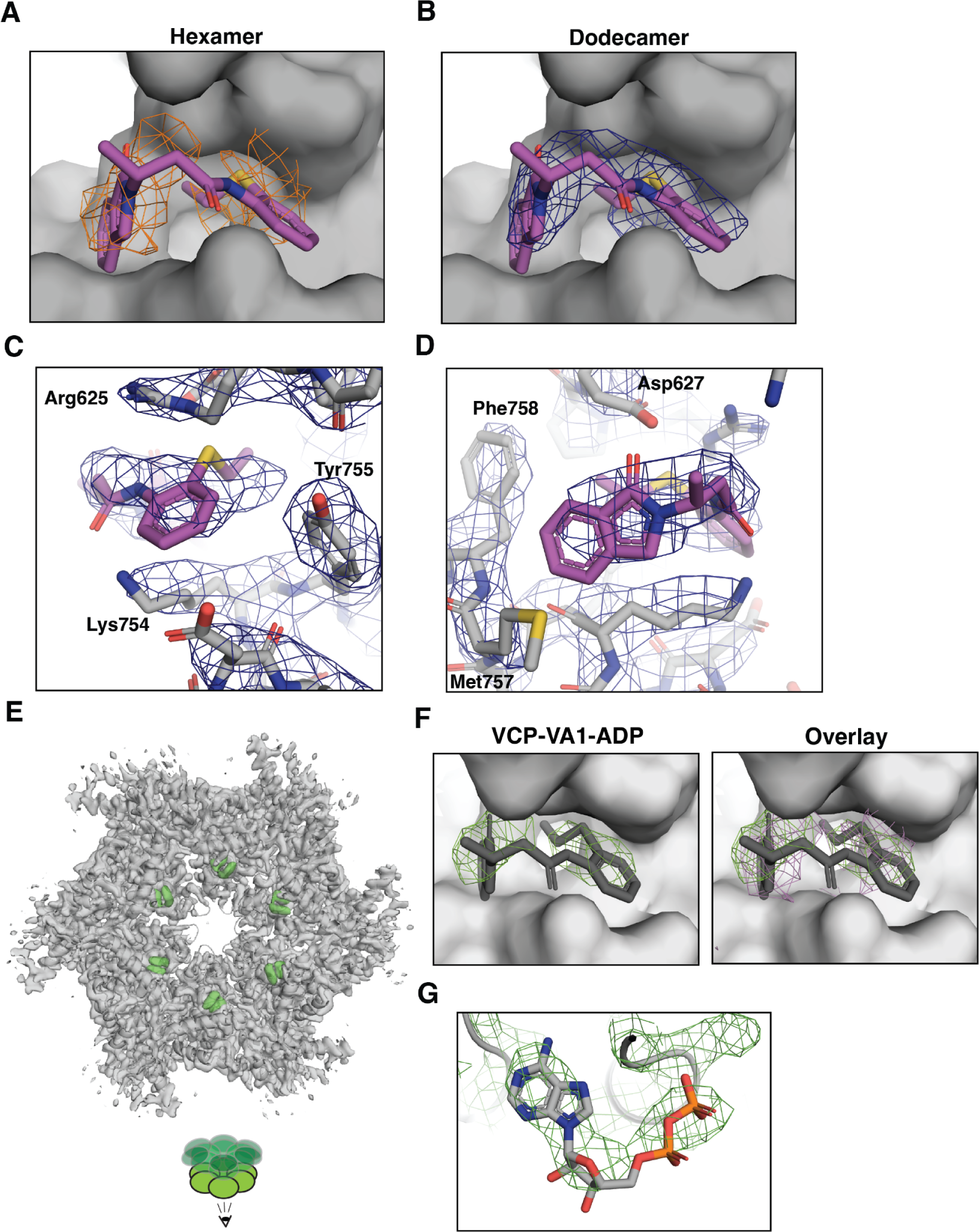
Structural analysis of VA1 binding VCP in two nucleotide states. (*A*) Model of VA1 in the proposed binding site (VA1, magenta, stick representation; VCP, gray, surface representation; VCP hexamer density, orange, mesh). (*B*) Model of VA1 in the proposed binding site in the dodecamer map (VA1, magenta, stick representation; VCP, gray, surface representation; VCP dodecamer density, blue, mesh). (*C*) View showing the fit of the VA1 thioether aniline (magenta, stick representation) as well as nearby residues (gray, stick representation) within the density map (mesh). (*D*) View showing the fit of the VA1 isoindoline (magenta, stick representation) as well as nearby residues (gray, stick representation) within the density map (mesh). (*E*) Molecular model of the VCP D2 domains (cartoon representation) in the VCP-VA1-ADP density map (semi-transparent surface with VA1 density highlighted in green). (*F*) Panels showing the VA1 binding site, comparing maps for datasets with and without the addition of ATP (VA1, gray, stick representation; VCP, gray, surface representation; VCP-VA1-ADP density, green, mesh; VCP-VA1-apo density, magenta, mesh). (*G*) Panel showing the D2 nucleotide binding site for VCP-VA1-ADP (ADP, gray, stick representation; VCP, gray, cartoon representation; density, green, mesh).

We next examined whether VA1 can bind to VCP in other nucleotide states in addition to the apo state. We incubated VCP with VA1 (50 µM), and subsequently added ATP (100 µM) before cryo-EM data collection (Fig. S3*B*, “VCP-VA1-ADP” dataset). 2D classification and 3D refinement (Fig. S3*B*) resulted in a map of the VCP hexamer D2 domains resolved to ∼3.8 Å (Fig. 4*E*). We carried out rigid body docking of individual domains of published VCP structures (PDB 5FTL for the D2 domains with C-terminal tails from PDB 7LN6)(10, 11) into the map, and made minor adjustments in the D2 domain near the VA1 binding site and in the nucleotide binding site (Fig. 4*E*). VA1 occupancy and orientation relative to nearby side chains is consistent between VCP-VA1-apo and VCP-VA1-ADP (Fig. 4*F*). We observed density corresponding to ADP, the hydrolyzed product of the added ATP, in the D2 active site, confirming a different nucleotide state than apo is compatible with VA1 binding (Fig. 4*G*). We additionally observed different states of pore loop II, which is proximal to the VA1 binding site, in our two structures, with a more stable pore loop in VCP-VA1-ADP than VCP-VA1-apo (Fig. S4*C*). Furthermore, the N-domain adopted a variety of conformations in our conditions (both with and without the addition of ATP) as indicated by 3D classification results in our cryo-EM processing (Fig. S4*D*). Together, our structural data indicate that VA1 binds a site near the VCP C-terminus in both apo and ADP-bound states of the D2 domain.

### The VA1 binding site allosterically regulates VCP activity

To probe the role of the VA1 binding site and test our compound binding model, we designed mutations in the C-terminal helix: 1) Tyr755-His, as a conservative mutation, and 2) Lys754-Asn, which is the amino acid at the equivalent position in the related AAA protein PEX1, a mutagenesis approach we have employed for other AAA proteins(36–38) (see Materials and Methods). We generated full-length recombinant VCP-K754N and VCP-Y755H and characterized these constructs using ATPase assays (Fig. S4*E-F*). The mutant constructs show K_1/2_ values similar to that of the WT protein, but, notably, different k_cat_ values than that of the WT protein (∼40% lower for VCP-K754N; ∼2-fold higher for VCP-Y755H) (VCP-K754N: K_1/2_ = 93 +/− 7 µM, k_cat_ = 0.051 +/− 0.015 s^-1^; VCP-Y755H: K_1/2_ = 83 +/− 5 µM, k_cat_ = 0.17 +/− 0.02 s^-1^; mean +/− s.d., N = 3) (Figs. 5*A-B* and S4*F*). Importantly, VA1 did not stimulate the ATPase activities of VCP-K754N or VCP-Y755H (Fig. 5*C*). We additionally assessed binding for VCP-Y755H using ITC, and observed no measurable binding of VA1 to VCP-Y755H (Fig. S4*G*).

**Fig. 5.**
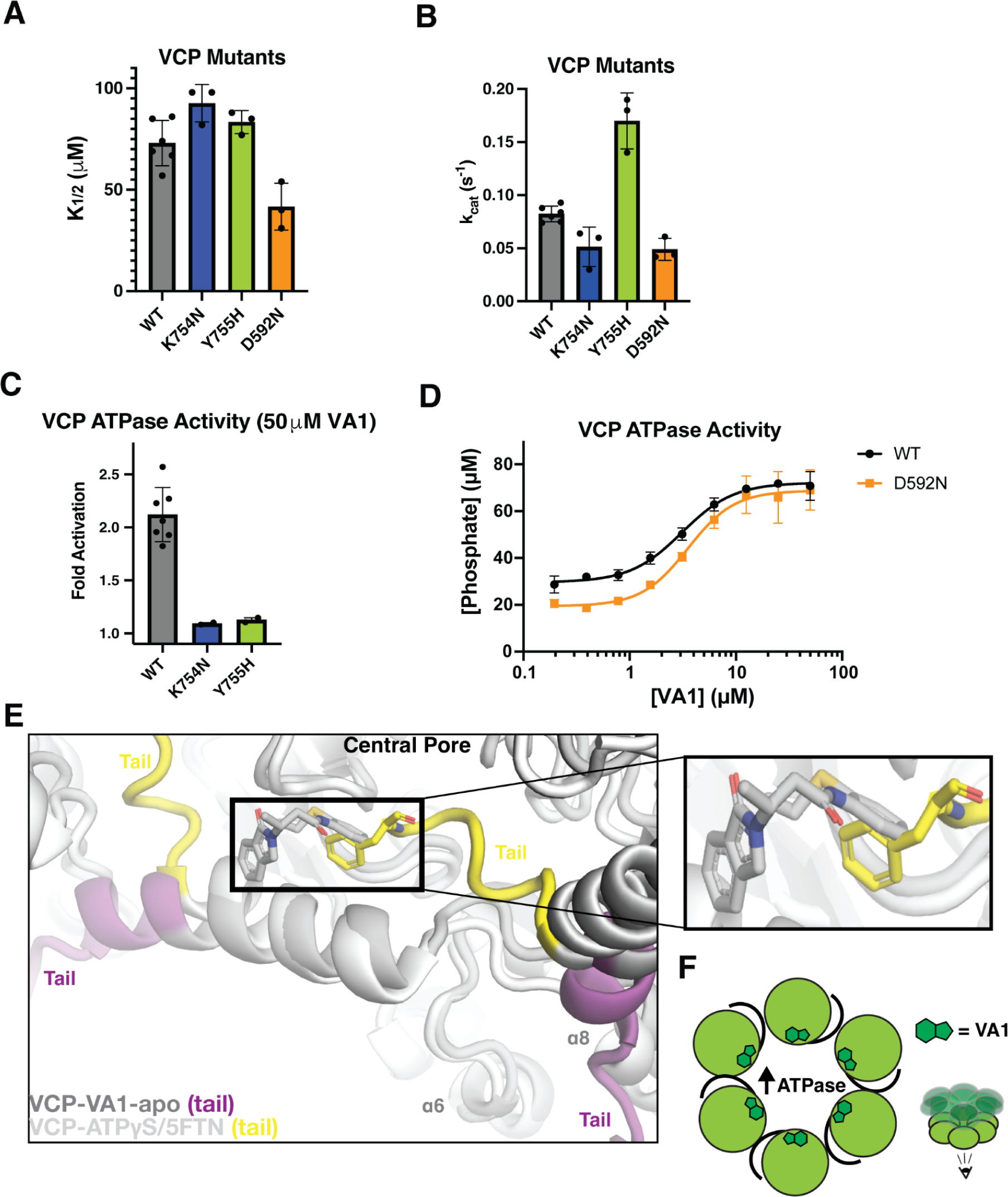
Characterization of the VA1 binding site. (*A-B*) Effect of the K754N, Y755H, and D592N mutations on VCP enzymatic activity. Graphs show values for the turnover number (K_1/2_, *A*) and ATP concentration required for half-maximal velocity (k_cat_, *B*) of VCP-WT, -K754N, -Y755H, and -D592N constructs, analyzed by measuring the steady-state ATPase rate across a range of ATP concentrations using a Malachite Green assay (mean +/− s.d., N = 6 independent experiments for WT; N = 3 independent experiments for mutants). (*C*) Fold change in ATPase activity of VCP (WT and K754N and Y755H mutants) in the presence of VA1 (50 µM ATP, 1 hr endpoint assay) (mean +/− s.d., N = 7 independent experiments for VCP-WT; mean +/− range, N = 2 independent experiments for VCP-K754N and VCP-Y755H). (*D*) Concentration-dependent activation of the ATPase activity of VCP (WT and D592N mutant) in the presence of VA1 (1 mM ATP, 1 hr endpoint assay) (mean +/− range, N = 2 independent experiments). (*E*) Alignment of VCP-VA1-apo (cartoon representation, gray, with C-terminal tails in magenta and VA1 in stick representation) and VCP-ATPγS (PDB: 5FTN) (cartoon representation, white, with C-terminal tails in yellow and Phe-768 in stick representation). Inset focuses on VA1. (*F*) Schematic for VA1 binding to VCP in a site that can be occupied by the C-terminal tail. VA1 accelerates ATP turnover by VCP.

To examine if VCP constructs with disease-associated mutations are activated by our compounds, we focused on Asp592-Asn, a mutation near the VA1 binding site that has been linked to ALS(39). We generated a recombinant construct of full-length VCP with the D592N mutation (see Materials and Methods) and characterized its ATP hydrolysis activity (Fig. S4*E-F*). The D592N mutation reduces both the K_1/2_ for ATP and the k_cat_ by ∼40% (K_1/2_ = 42 +/− 9 µM, k_cat_ = 0.049 +/− 0.009 s^-1^, mean +/− s.d., N = 3) (Figs. 5*A-B* and S4*F*). However, VA1 treatment restores the activity of VCP-D592N, with an EC_50_ comparable to the WT protein (EC_50_ of 3.5 +/− 0.4 µM, mean +/− range, N = 2, 1 mM ATP) (Fig. 5*D*). Our data suggest that resistance-conferring mutations in the VA1 binding site can modulate ATP turnover by VCP, and while a disease mutation proximal to the binding site reduces enzymatic activity, it does not affect VA1-mediated activation.

## Discussion

Here, we report the discovery of an allosteric site in VCP that binds a small molecule activator. We identified VA1, a small molecule that stimulates VCP activity, using an unbiased screen, and carried out cryo-EM studies of VCP in complex with VA1 in ADP-bound and apo states of the D2 active site. Engineered mutations in the VA1 binding pocket confer resistance to VA1 and result in both increases and decreases in VCP ATP turnover, suggesting the sensitivity of the enzyme to modulation of this allosteric site.

Our structural data suggest how VCP is allosterically activated by VA1 binding in the D2 domain. We analyzed the VA1 binding site in available structures of the VCP hexamer. In our VA1-bound structures, the C-terminal tail is positioned between helices ɑ6 and ɑ8 of the neighboring D2 domain (Fig. 5*E*). In contrast, in previously solved structures of VCP in an ATPγS-bound D2 state, the C-terminal tail orients toward the neighboring D2 nucleotide binding site(10, 35, 40) (Fig. 5*E*). In this conformation, Arg-766 coordinates the ATPγS γ-phosphate, and, interestingly, Phe-768 orients into the VA1 binding pocket (Figs. 5*E* and S4*H*). VA1 binding may structurally mimic the interactions made by Phe-768, impacting conformational changes in and around the pocket. Further, the displacement of the C-terminal tail may change the interactions made by this motif to enhance enzymatic activity (Fig. 5*F*). Consistent with the importance of this site for regulation of VCP activity, we observed changes in turnover number (k_cat_) with the introduction of the K754N and Y755H mutations (Fig. 5*B*). The C-terminal tail has been proposed to contain at least three phosphorylation sites (Thr-761, Ser-784, Tyr-805)(19, 41–44). Additionally, a subset of VCP cofactors, including UBXD1, PLAA, and PNGase, have been proposed to bind the C-terminal tail(16, 45–48). Together, these data suggest that VCP’s C-terminal tail and the motifs with which it interacts in VCP form an important regulatory site that can be targeted by small molecule activators.

Accumulation of toxic protein aggregates is a common feature of degenerative diseases(49). VCP disassembles aggregated proteins *in vitro* and in cells, suggesting its activation as a means of removing disease aggregates(7). Further to this rationale for activating wild-type VCP, mutations in VCP have been linked to degenerative diseases by multiple lines of evidence, including cell biological studies, mouse models, and patient sequencing data(6, 7). Loss of VCP ATPase activity can result from mutations such as the D395G mutation associated with vacuolar tauopathy and the ALS-associated D592N mutation characterized herein(7). Other mutations result in no change to or an increase in VCP’s ATPase activity, but the activity level does not correlate with disease severity, and, importantly, disease phenotypes are better imitated by loss-of-function models(18, 50). The mutations may cause an altered equilibrium between conformational states of the N-terminal domain in cells, consistent with measured changes in VCP’s affinity for certain cofactors (e.g., the Ufd1/Npl4 complex and UBXD1)(18, 51, 52). Testing differential binding of cofactors in the presence of chemical activators can help dissect this further, and examine VCP’s potential to clear disease aggregates in general. At this stage, the micromolar potency of VA1 may limit its measurable on-target activity in cells. However, with our structural data and mechanistic hypothesis for activation, structure-based drug design campaigns will lead to more potent compounds.

Protein cofactors known to stimulate VCP ATPase activity bind near the N-terminus(11, 14, 15, 25), but the dispersed surface contacts involved in protein-protein interactions may not be readily recapitulated by small molecule binders. The VA1 binding site is distinct from this N-terminal regulatory site and suggests a unique mechanism of activation that may be compatible with multiple interacting partners. We note that the primary structural elements that comprise the VA1 binding site, namely, the loop containing Arg-625/Asp-627 and the C-terminal helix, are conserved in a subset of AAA proteins, and could function as a regulatory site for these additional mechanoenzymes. Interestingly, it is possible that the VA1 binding pocket could serve as a binding site for naturally occurring metabolites that regulate AAA proteins. The structures of VCP in complex with VA1 open the door to rational structure-based design of activators targeting an allosteric regulatory site in this important AAA protein central to cellular proteostasis.

## Materials and Methods

### Plasmids

Plasmid for expressing wild-type *Homo sapiens* VCP was obtained from T.-F. Chou (Caltech). Plasmids for VCP mutants were generated by site-directed mutagenesis (including VCP-ND1L, which was generated by mutating Gln-473 to a stop codon).

### Expression and purification of VCP constructs

The VCP constructs were expressed in *E. coli* Rosetta (DE3) pLysS cells (Merck, cat. No. 70954) grown in Miller’s LB medium (LMM, Formedium, cat. No. LMM105). Culture growth at 37 °C was monitored by measuring absorbance at 600 nm (A_600_) and protein expression was induced at A_600_ = ∼0.7 with 0.5 mM IPTG (GoldBio). The culture was grown at 18 °C for ∼15 hr, pelleted, and resuspended in lysis buffer (50 mM K.HEPES pH 7.5, 400 mM KCl, 20 mM imidazole, 5% glycerol, 5 mM β-mercaptoethanol, 200 µM MgATP, 1 mM *p*-phenylmethylsulfonyl fluoride, cOmplete EDTA-free protease inhibitor cocktail (Roche)). All subsequent purification steps were performed at 4 °C. Cells were lysed using an EmulsiFlex C5 homogenizer (Avestin, ∼6 cycles at 10,000-15,000 psi homogenization pressure). The lysate was clarified by centrifugation at 45,000 rpm for 30 min using a Type 70 Ti rotor in a Beckman Coulter Optima LE-80K ultracentrifuge.

The clarified lysate was loaded onto Ni-NTA resin and incubated for 1-2 hr. The resin was washed with ∼200 mL of wash buffer (50 mM K.HEPES pH 7.5, 400 mM KCl, 20 mM imidazole, 5% glycerol, 5 mM β-mercaptoethanol, 200 µM MgATP) and eluted with elution buffer (50 mM K.HEPES pH 7.5, 400 mM KCl, 300 mM imidazole, 5% glycerol, 5 mM β-mercaptoethanol, 100 µM MgATP). The eluate was treated with tobacco etch virus (TEV) protease (∼2 mg) and dialyzed in dialysis buffer (50 mM K.HEPES pH 7.5, 100 mM KCl, 20 mM imidazole, 1 mM MgCl_2_, 5% glycerol, 5 mM β-mercaptoethanol, 100 µM MgATP) overnight. Protein was passed 3 times over Ni-NTA resin pre-equilibrated with reverse Ni buffer (25 mM K.HEPES pH 8, 100 mM KCl, 20 mM imidazole, 1 mM MgCl_2_, 5% glycerol, 5 mM β-mercaptoethanol), or incubated 1 hr then eluted. Protein was loaded onto a Q column (GE Healthcare) pre-equilibrated with low salt buffer (25 mM K.HEPES pH 8, 100 mM KCl, 1 mM MgCl_2_, 5% glycerol, 2 mM DTT) and eluted in a 0-100% gradient of high salt buffer (50 mM K.HEPES pH 8, 500 mM KCl, 1 mM MgCl_2_, 5% glycerol, 2 mM DTT) over 40 mL. Fractions containing VCP were concentrated using an Amicon Ultra 50K concentrator to 1-2 mL, filtered, and loaded into a Superdex 200 16/600 column (GE Healthcare) equilibrated with gel filtration buffer (50 mM K.HEPES pH 7.5, 200-250 mM KCl, 1-2 mM MgCl_2_, 5% glycerol, 1 mM reducing agent (DTT or TCEP)). The fractions containing purified VCP were pooled, concentrated using an Amicon Ultra 50K concentrator, and frozen in liquid nitrogen and stored at −80 °C.

### Mass photometry

Data were collected using a OneMP mass photometer (Refeyn) calibrated with bovine serum albumin (ThermoFisher, cat. No. 23210), beta amylase (Sigma Aldrich, cat. No. A8781-1VL), and thyroglobulin (Sigma Aldrich, cat. No. T9145-1VL). Focus was adjusted using filtered (0.22 µm) phosphate-buffered saline (PBS), then a 400 nM solution of VCP in filtered PBS with 2% DMSO was added directly, diluting it 4-fold. Movies were acquired for 6,000 frames (60 s) using AcquireMP software (version 2.4.0) and default settings. Raw data were converted to frequency distributions using DiscoverMP software (10 kDa bin size).

### Analysis of ATPase activity

The ATPase activity of VCP was examined using Malachite Green and ADPGlo assays. For Malachite Green assays to determine kinetic parameters, incubation of VCP with ATP proceeded for different time points before simultaneous addition of reagent to read out phosphate production as absorbance at 620 nm using a Synergy NEO microplate reader. The rates from control reactions without VCP were subtracted from the corresponding reactions with VCP at the same [ATP]. VCP was assayed at 100-200 nM in 50 mM K.HEPES pH 7.5, 25 mM KCl, 2.5 mM MgCl_2_, 2.5 mM GSH, 0.01% Triton-X-100, 0.1 mg/mL BSA. Analyses of activators were carried out in the same conditions, with a 5 min pre-incubation of VCP with compound before addition of ATP, and a single 60 min time point of enzymatic activity. The final DMSO concentration was 0.5%. ADPGlo assays to test activators were carried out under the same ATPase reaction conditions as Malachite Green assays to test activators, except using 10-100 nM VCP. After reagent incubations, generation of ADP was measured as luminescence using a Synergy NEO microplate reader.

In line with our previous work on AAA proteins, we analyzed the kinetic parameters of VCP using a modified Michaelis-Menten equation, which includes a Hill coefficient(36–38):

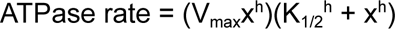

Where V_max_ is the maximum ATPase rate, h is the Hill coefficient, x is ATP concentration, and K_1/2_ is the concentration of ATP required for half-maximal enzyme rate. The K_1/2_ term distinguishes this constant from the standard Michaelis-Menten constant K_m_. Catalytic turnover number (k_cat_) was calculated by dividing V_max_ by the concentration of VCP in the assay.

For the compound EC_50_ determination, for each experiment the fold-change in enzyme activity in the presence of compound compared to control was plotted against compound concentrations, and the data were fit using a sigmoidal dose-response curve equation:

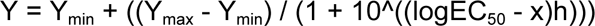

### Small molecule activator screening protocol

The Z’ of the ADPGlo assay with conditions used for screening was 0.86. The screening campaign was performed against a ∼35,000-compound library in a 384-well format. After a 5 min preincubation of 25 nM VCP with 20 µM compound, ATP was added to 100 µM and the reaction was allowed to proceed for 60 min (∼10-15% ATP consumption in controls) before addition of ADPGlo reagent (40 min incubation) and Kinase Detection Reagent (40 min incubation). The hit rate using a 2x s.d. (∼20% activation) cutoff was ∼0.21%. Primary hits that repeated with the same experimental setup, but did not result in increased luminescence in the absence of VCP, were tested for dose-responsiveness. Dose-responsive hits with favorable physicochemical and structural characteristics were re-purchased from the supplier for re-testing, and Compound **1** was selected as the highest potency compound.

### Chemical synthesis and characterization

#### Reagent Abbreviations

DCM: dichloromethane

DMF: dimethyl formamide

HOAc: acetic acid

MeCN: acetonitrile

NMI: N-methylimidazole

TCFH: Chloro-N,N,N’,N’-tetramethylformamidinium hexafluorophosphate

THF: tetrahydrofuran

**Compound 1 (hit re-synthesis).** To a stirred solution of precursor **1** (1 eq.) in DCM (0.3 M), was added oxalyl chloride (1.5 eq.), followed by one drop of DMF as catalyst. The reaction was stirred for ∼30 mins; the acyl chloride **2** was pure enough and could be used without further purification. Aniline **3** (1 eq.) and THF (V_DCM_:V_THF_ = 4:1) were added, followed by addition of triethylamine (2 ∼ 4 eq.), then the reaction was stirred at room temperature. When the reaction completed (as monitored by LC/MS), the product was purified by silica gel chromatography with hexane/ethyl acetate to afford product QL1032 (Compound **1**) as white powder, 83% yield.

^1^H NMR (600 MHz, Chloroform-*d*) δ 8.54 (s, 1H), 8.34 (d, J = 8.2 Hz, 1H), 7.84 (dd, J = 7.6, 1.1 Hz, 1H), 7.51 (tt, J = 7.5, 1.0 Hz, 1H), 7.49 – 7.45 (m, 1H), 7.44 (d, J = 7.6 Hz, 1H), 7.41 (dt, J = 7.6, 0.9 Hz, 1H), 7.35 – 7.28 (m, 1H), 7.08 – 7.01 (m, 1H), 4.52 (s, 2H), 4.02 (t, J = 6.3 Hz, 2H), 2.89 (t, J = 6.3 Hz, 2H), 2.66 (q, J = 7.3 Hz, 2H), 1.10 (t, J = 7.3 Hz, 3H).

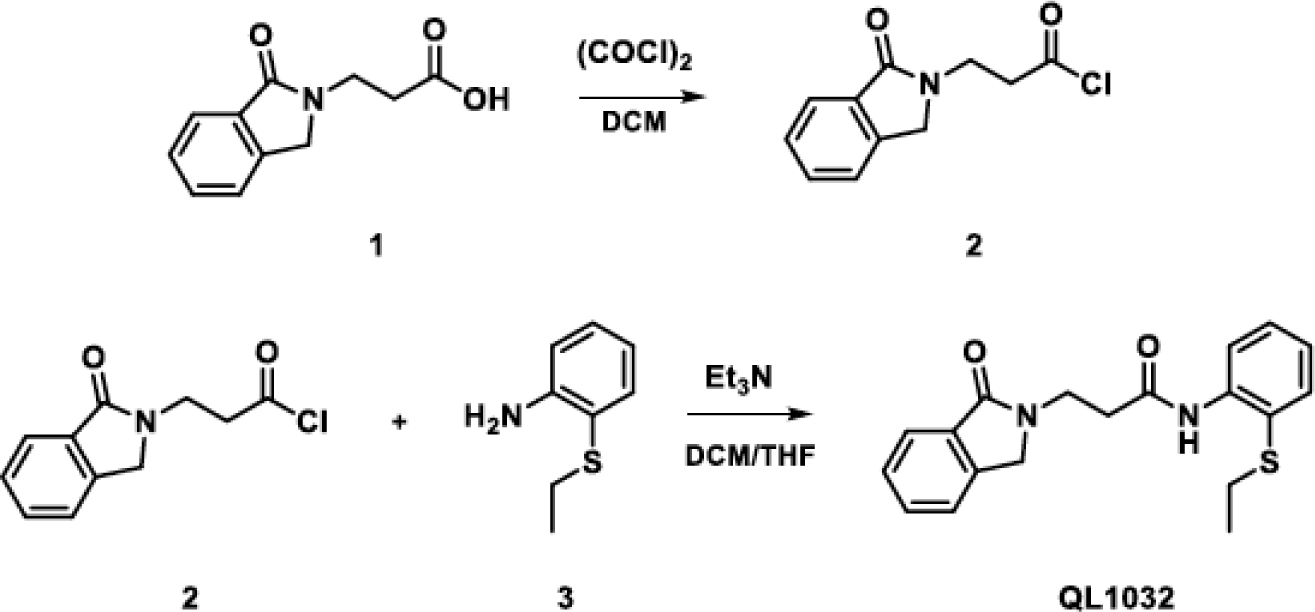

Reagents: 2-(Ethylsulfanyl)aniline, purity 95%, Enamine EN300-49598

3-(1-oxo-2,3-dihydro-1H-isoindol-2-yl)propanoic acid, purity 95%, Enamine EN300-186584

**Compound 2.** Same general method as for Compound **1**. Product was white powder, 53% yield.

^1^H NMR (600 MHz, Chloroform-*d*) δ 8.34 (d, *J* = 8.0 Hz, 1H), 7.93 – 7.76 (m, 2H), 7.51 (t, *J* = 7.4 Hz, 1H), 7.44 (t, *J* = 7.5 Hz, 1H), 7.41 (d, *J* = 7.6 Hz, 1H), 7.01 (t, *J* = 7.8 Hz, 1H), 6.93 (t, *J* = 7.8 Hz, 1H), 6.83 (d, *J* = 8.1 Hz, 1H), 4.52 (s, 2H), 4.03 (dt, *J* = 18.5, 6.6 Hz, 4H), 2.85 (t, *J* = 6.3 Hz, 2H), 1.38 (t, *J* = 7.0 Hz, 3H).

Reagents: 2-Ethoxyaniline, purity >98%, TCI #P0089

3-(1-oxo-2,3-dihydro-1H-isoindol-2-yl)propanoic acid, purity 95%, Enamine, EN300-186584

**Compound 3.** General method scheme shown below. Step 1: Carboxylic acid **1** was synthesized by reported method(54) from *O*-Phthalaldehyde and *β*-homo-amino acid; the crude can be used without further purification. Step 2: To a stirred solution of carboxylic acid **1** (1 eq.) in DCM (0.3 M), was added oxalyl chloride (1.5 eq.), followed by one drop of DMF as catalyst, the reaction was stirred for ∼30 mins. The acyl chloride **4** was pure enough and could be used without further purification. Aniline **2** (1 eq.) and THF (V_DCM_:V_THF_ = 4:1) was added, followed by addition of triethylamine (2 - 4 eq.), then the reaction was stirred at room temperature. When reaction was completed (monitored by LC/MS), purified by silica gel chromatography with hexane/ethyl acetate to afford Compound **3** as white powder, 88% yield.

^1^H NMR (600 MHz, Chloroform-*d*) δ 8.61 (s, 1H), 8.25 (d, *J* = 8.2 Hz, 1H), 7.81 (dd, *J* = 7.3, 1.3 Hz, 1H), 7.51 (td, *J* = 7.4, 1.2 Hz, 1H), 7.46 – 7.44 (m, 1H), 7.43 – 7.41 (m, 2H), 7.26 (ddd, *J* = 8.4, 5.9, 1.6 Hz, 1H), 7.02 (td, *J* = 7.5, 1.4 Hz, 1H), 4.69 (dt, *J* = 7.9, 6.5 Hz, 1H), 4.48 (s, 2H), 3.15 (dd, *J* = 14.7, 8.1 Hz, 1H), 2.80 (dd, *J* = 14.7, 6.1 Hz, 1H), 2.68 (dd, *J* = 7.3, 3.0 Hz, 2H), 1.53 (d, *J* = 6.9 Hz, 3H), 1.14 (t, *J* = 7.3 Hz, 3H).

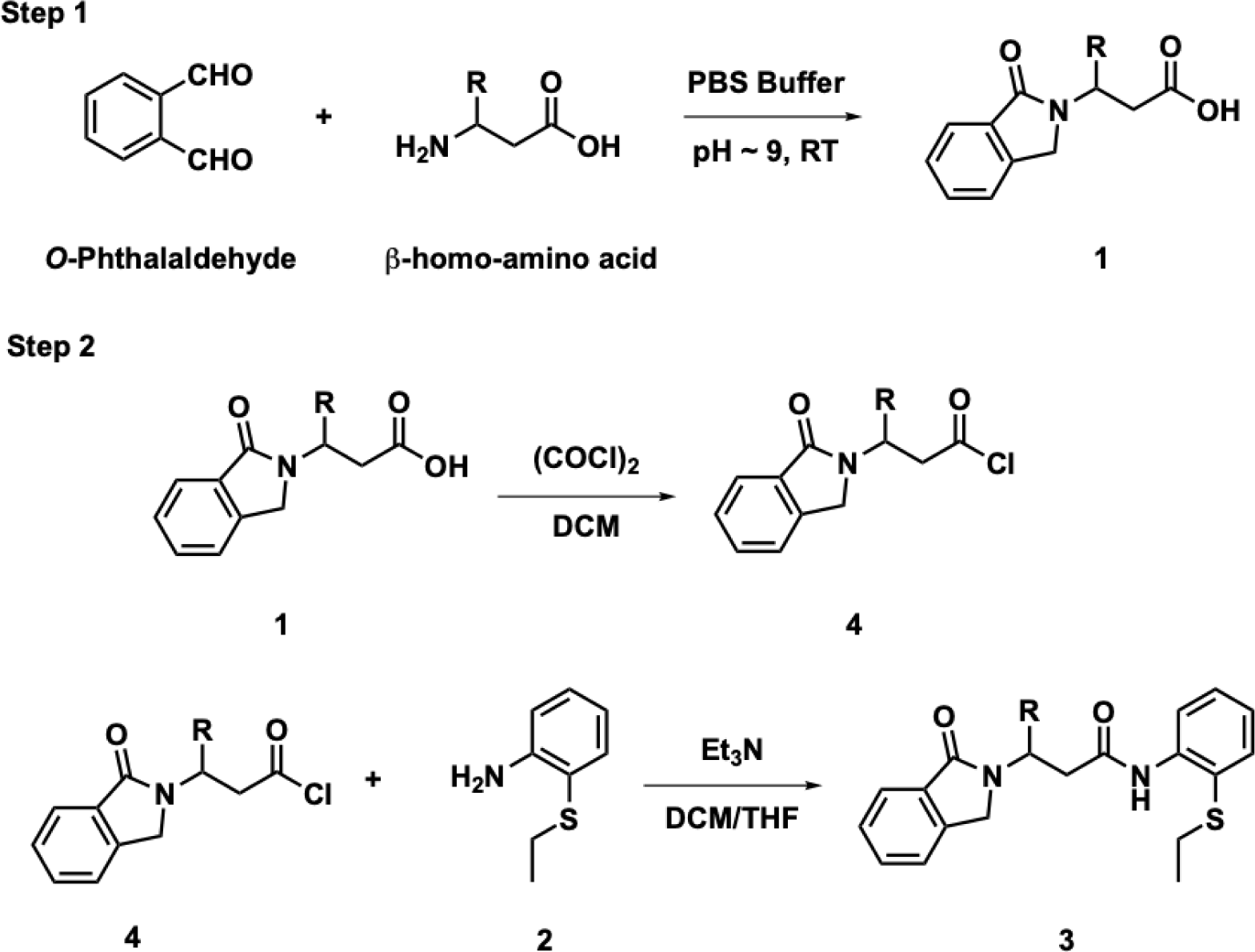

Reagents: 2-(Ethylsulfanyl)aniline, purity 95%, Combi-Blocks QA-0283

*O*-Phthalaldehyde, purity 97%, Combi-Blocks SS-7380

DL-3-aminobutyric acid, purity 96%, Combi-Blocks QB-7615

**Compound 4.** Same general method as for compound **3**. Product was a white powder (19% yield).

^1^H NMR (600 MHz, Chloroform-*d*) δ 8.53 (s, 1H), 8.17 (d, *J* = 8.2 Hz, 1H), 7.77 (d, *J* = 7.4 Hz, 1H), 7.49 (td, *J* = 7.5, 1.2 Hz, 1H), 7.41 (dd, *J* = 7.7, 1.4 Hz, 2H), 7.39 (d, *J* = 7.5 Hz, 1H), 7.22 (ddd, *J* = 8.6, 7.4, 1.6 Hz, 1H), 6.99 (td, *J* = 7.6, 1.3 Hz, 1H), 4.51 (s, 2H), 4.02 (td, *J* = 10.2, 4.0 Hz, 1H), 3.28 (dd, *J* = 15.0, 10.5 Hz, 1H), 2.92 (dd, *J* = 15.0, 4.0 Hz, 1H), 2.71 – 2.58 (m, 2H), 2.46 (dp, *J* = 9.7, 6.7 Hz, 1H), 1.10 (dd, *J* = 8.5, 6.9 Hz, 6H), 0.94 (d, *J* = 6.7 Hz, 3H).

Reagents: 2-(Ethylsulfanyl)aniline, purity 95%, Combi-Blocks QA-0283

*O*-Phthalaldehyde, purity 97%, Combi-Blocks SS-7380

Boc-L-beta-homovaline, purity 98%, Combi-Blocks SS-1159

**Compound 5.** Step 1: Carboxylic acid **1** was synthesized by reported method(54) from *O*-Phthalaldehyde and *β*-homo-amino acid; the crude can be used without further purification. Step 2: The carboxylic acid **1** (1.0 equiv), aniline **2** (1.2 equiv), and NMI (2.1 equiv) were combined and dissolved in MeCN (0.2 M); TCFH (1.1 equiv) was added in a single portion(55). The reaction stirred at room temperature (monitored by LC/MS). The reaction was then concentrated before purification by silica gel chromatography with hexane/ethyl acetate to afford Compound **5** as white powder (69% yield).

^1^H NMR (600 MHz, Chloroform-*d*) δ 8.54 (s, 1H), 8.16 (d, *J* = 8.3 Hz, 1H), 7.77 (dt, *J* = 7.6, 1.0 Hz, 1H), 7.49 (td, *J* = 7.5, 1.2 Hz, 1H), 7.43 – 7.40 (m, 2H), 7.40 – 7.37 (m, 1H), 7.22 (ddd, *J* = 8.5, 7.5, 1.6 Hz, 1H), 7.02 – 6.96 (m, 1H), 4.51 (s, 2H), 4.02 (td, *J* = 10.2, 4.0 Hz, 1H), 3.28 (dd, *J* = 15.0, 10.5 Hz, 1H), 2.92 (dd, *J* = 15.0, 4.0 Hz, 1H), 2.72 – 2.56 (m, 2H), 2.46 (dp, *J* = 9.8, 6.7 Hz, 1H), 1.10 (dd, *J* = 8.2, 6.7 Hz, 6H), 0.94 (d, *J* = 6.7 Hz, 3H).

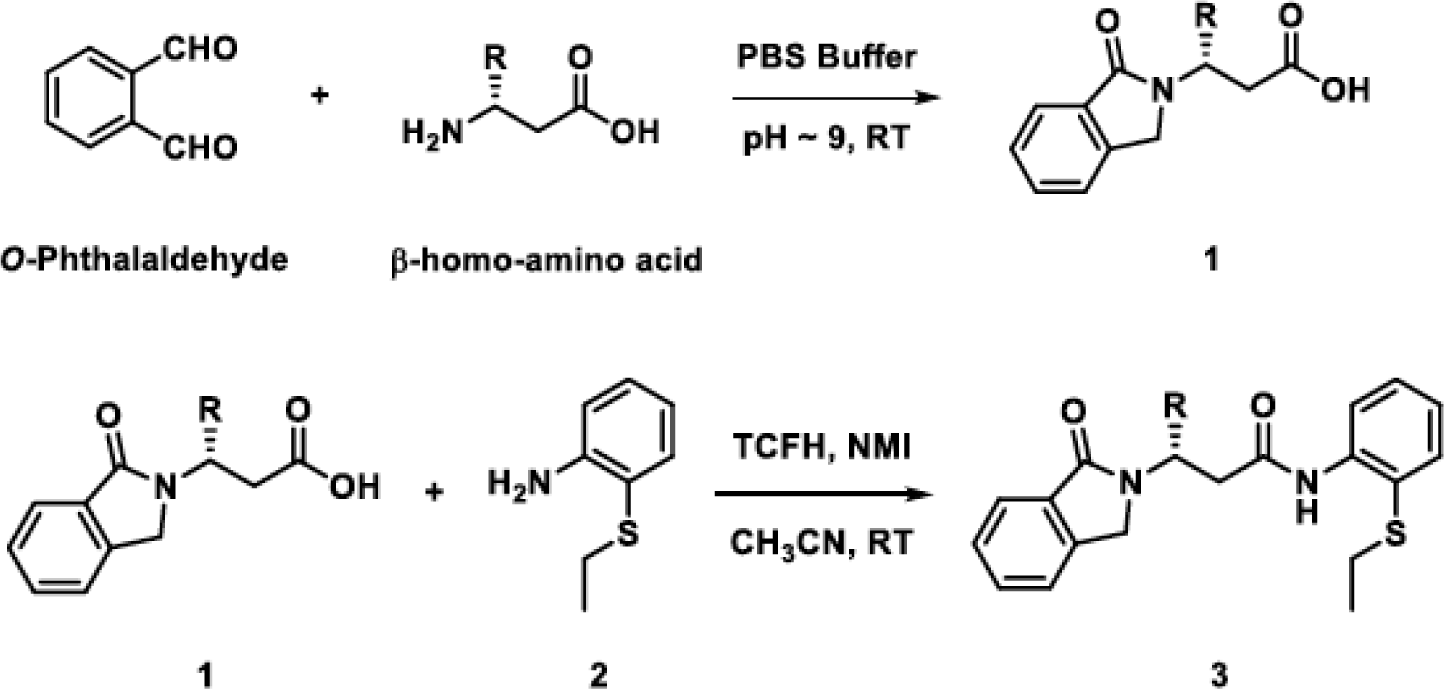

Reagents: 2-(Ethylsulfanyl)aniline, purity 95%, Enamine EN300-49598

*O*-Phthalaldehyde, purity 97%, Combi-Blocks SS-7380

(*S*)-3-Amino-4-methyl-pentanoic acid, purity 98%, Combi-Blocks OR-9452

**Compound 6 (VA1).** Step 1: Carboxylic acid **1** was synthesized by reported method(54) from *O*-Phthalaldehyde and *β*-homo-amino acid; the crude can be used without further purification. Step 2: The carboxylic acid **1** (1.0 equiv), 2-(ethylsulfanyl)aniline (**2**) (1.2 equiv), and NMI (2.1 equiv) were combined and dissolved in MeCN (0.2 M); TCFH (1.1 equiv) was added in a single portion(55). The reaction stirred at room temperature (monitored by TLC). The reaction was then concentrated before purification by silica gel chromatography with hexane/ethyl acetate to give product **3** (Compound 6, VA1) as a white powder (80% yield).

1H NMR (600 MHz, Chloroform-*d*) δ 8.60 (s, 1H), 8.27 (d, *J* = 8.2 Hz, 1H), 7.81 (d, *J* = 7.5 Hz, 1H), 7.51 (t, *J* = 7.5 Hz, 1H), 7.45 (d, *J* = 4.1 Hz, 1H), 7.43 (m, 2H), 7.30 – 7.26 (m, 1H), 7.02 (t, *J* = 7.6 Hz, 1H), 4.68 (q, *J* = 7.0 Hz, 1H), 4.49 (s, 2H), 3.16 (dd, *J* = 14.7, 8.1 Hz, 1H), 2.80 (dd, *J* = 14.7, 6.1 Hz, 1H), 2.68 (qd, *J* = 7.3, 2.9 Hz, 2H), 1.53 (d, *J* = 6.9 Hz, 3H), 1.14 (t, *J* = 7.3 Hz, 3H).

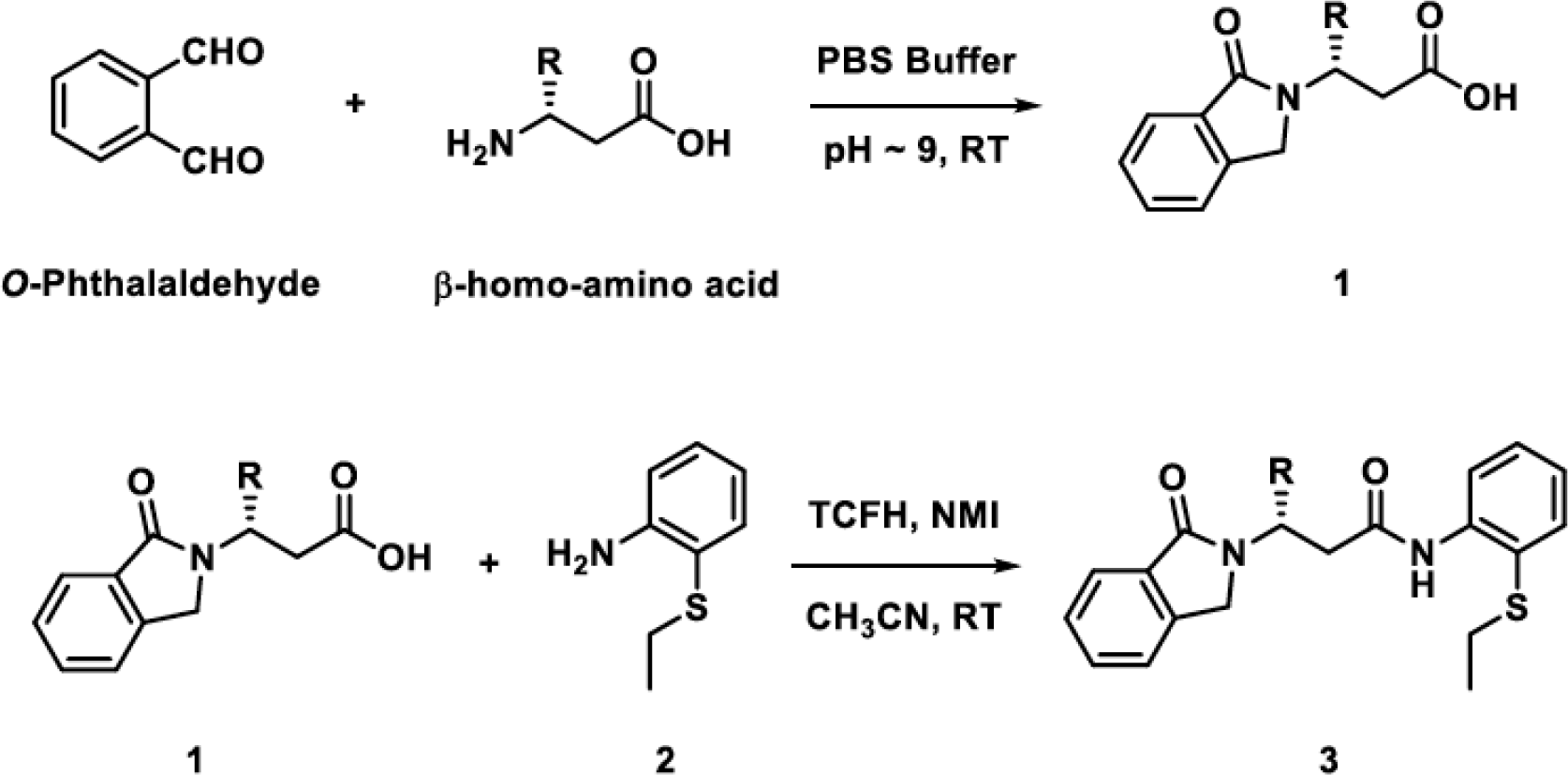

Reagents: 2-(Ethylsulfanyl)aniline, purity 95%, Enamine EN300-49598

*O*-Phthalaldehyde, purity 97%, Combi-Blocks SS-7380

(*R*)-3-Aminobutanoic acid, purity 97%, AmBeed A226851-005

### Isothermal titration calorimetry

ITC measurements were performed using a MicroCal auto-iTC200 calorimeter (MicroCal). Purified VCP constructs were diluted to 30 µM in 50 mM K.HEPES pH 7.5, 25 mM KCl, 2.5 mM MgCl_2_, 1 mM TCEP, 0.01% Triton-X-100, and DMSO was added to 0.6%. A separate buffer solution was prepared to match the final concentrations of components accounting for the contribution of the buffer the VCP was stored in. Final assay buffer concentrations were 50 mM K.HEPES pH 7.5, 30-64 mM KCl, 2.5 mM MgCl_2_, 0-0.1 mM DTT, 1 mM TCEP, 0.1-1.1% glycerol, 0.008-0.01% Triton-X-100. DMSO solutions of compounds were diluted in the assay buffer to 30 µM (0.6% DMSO, final concentration) and titrated into the diluted VCP in the ITC chamber (25 °C, stirring at 750 rpm). Titrations were carried out with 19 injections of 2 µL, with a duration of 4 s per injection and spacing of 150 s. Independent titrations of the same compound solutions into buffer (with 0.6% DMSO, without VCP) were carried out and subtracted from the data for the titrations into VCP. The data were analyzed using MicroCal PEAQ-ITC software and plotted using Prism (version 9.2, GraphPad Software). Dissociation constants from each titration of VA1 into full-length VCP-WT were obtained using a single-site equilibrium-binding model, carried out in the MicroCal PEAQ-ITC software.

### Cryo-EM sample preparation

Full-length VCP-WT was buffer exchanged into 50 mM K.HEPES pH 7.5, 25 mM KCl, 2.5 mM MgCl_2_, 2.5 mM GSH using an Amicon Ultra 100K concentrator, and concentrated to ∼1 mg/mL. For the dataset without nucleotide (VCP-VA1-apo), VA1 (200 µM, 0.5% v/v DMSO, final concentrations) was added and incubated at room temperature 5 min, then fluorinated octyl maltoside (FOM) detergent was added (0.01% (w/v) final concentration). Sample was applied to Graphene Oxide on Quantifoil R2/2 300-square-mesh copper grids. Samples were incubated on grids on a FEI Vitrobot IV for 30 s and blotted for 3.5-4 s at 100% humidity and 25 °C (force: 0, wait time: 0 s), then plunged into liquid ethane.

For the dataset with addition of nucleotide (VCP-VA1-ADP), VA1 (50 µM, 0.5% DMSO, final concentrations) was added and the sample incubated at room temperature 5 min. MgATP (100 µM, final concentration) was subsequently added to the VCP/VA1 sample, and incubated at room temperature 5 min, then fluorinated octyl maltoside (FOM) detergent was added (0.01% (w/v) final concentration). Sample was applied to freshly glow-discharged (30 s) Quantifoil R2/2 300-square-mesh copper grids and blotted for 3.5-4 s at 100% humidity and 25 °C (force: 0, wait time: 0 s) on a FEI Vitrobot IV, then plunged into liquid ethane.

### Cryo-EM data collection

Micrographs were recorded using automated data collection (SerialEM or Leginon)(56, 57) on a FEI Titan Krios with a Gatan K2 or K3 camera in super-resolution mode. Collection settings (acceleration voltage, camera, CS corrector, energy filter, exposure time, frame number, pixel size, dose) are listed in Table S2.

### Cryo-EM data processing

Correction of inter-frame movement for each pixel and dose-weighting was performed using MotionCor2 or Relion’s own implementation(58–60). CTF parameters were estimated using CTFFIND4(61). Further processing was carried out using Relion v.3.1 or v.4.0(59, 60). For VCP-VA1-apo, a small set of particles was manually picked and subjected to 2D classification to generate references for automated particle picking. Autopicked particles were cleaned with multiple rounds of 2D classification, and subsequently selected hexamer and dodecamer classes. Cleaned hexamer particles were subjected to 3D classification to select higher quality and well-balanced subsets of particles. These particles were subjected to global 3D refinement without symmetry to yield an overall VCP reconstruction. To improve density near the proposed VA1 binding site, we generated a mask for the D2 domains and performed signal subtraction to remove N-terminal domain and D1 domain signal from raw particle images. Subtracted particles were subjected to local 3D refinement with C6 symmetry. Other datasets were similarly processed with minor modifications (Fig. S3).

### Cryo-EM model building

A VCP model (5FTL) was segmented into N-terminal, D1, and D2 domains for each chain, and coordinates for the C-terminal tail were segmented from another VCP model (7LN6). Each segment was rigid body fitted in Chimera(62) and Coot (“jiggle fit” function)(63). These docked domains were joined to generate the overall VCP hexamer model. Real space sphere refinement in Coot was used to adjust side chains in the VA1 binding pocket and the D2 nucleotide binding pocket.

## Acknowledgements

We thank T.-F. Chou for providing the *H. sapiens* VCP plasmid. T.M.K. is grateful to the NIH (GM130234) and Starr Cancer Consortium (I12-0055) for supporting this research. N.H.J. was supported in part by the National Science Foundation Graduate Research Fellowship Program (2017242069). L.E.V. was supported in part by the NIH T32 GM115327 and GM136640 Chemistry-Biology Interface Training Grant to the Tri-Institutional PhD Program in Chemical biology. We are grateful to the Fisher Drug Discovery Resource Center at The Rockefeller University for assistance with the high-throughput screen and instrument use. We also thank M. Ebrahim, J. Sotiris, and H. Ng, the Evelyn Gruss Lipper Cryo-Electron Microscopy Resource Center, and the New York Structural Biology Center for Cryo-EM support. This work was supported in part by The Rockefeller University Robertson Therapeutic Development Fund and The Rockefeller University Kellen Women’s Entrepreneurship Fund.

---

**Fig. S1.**
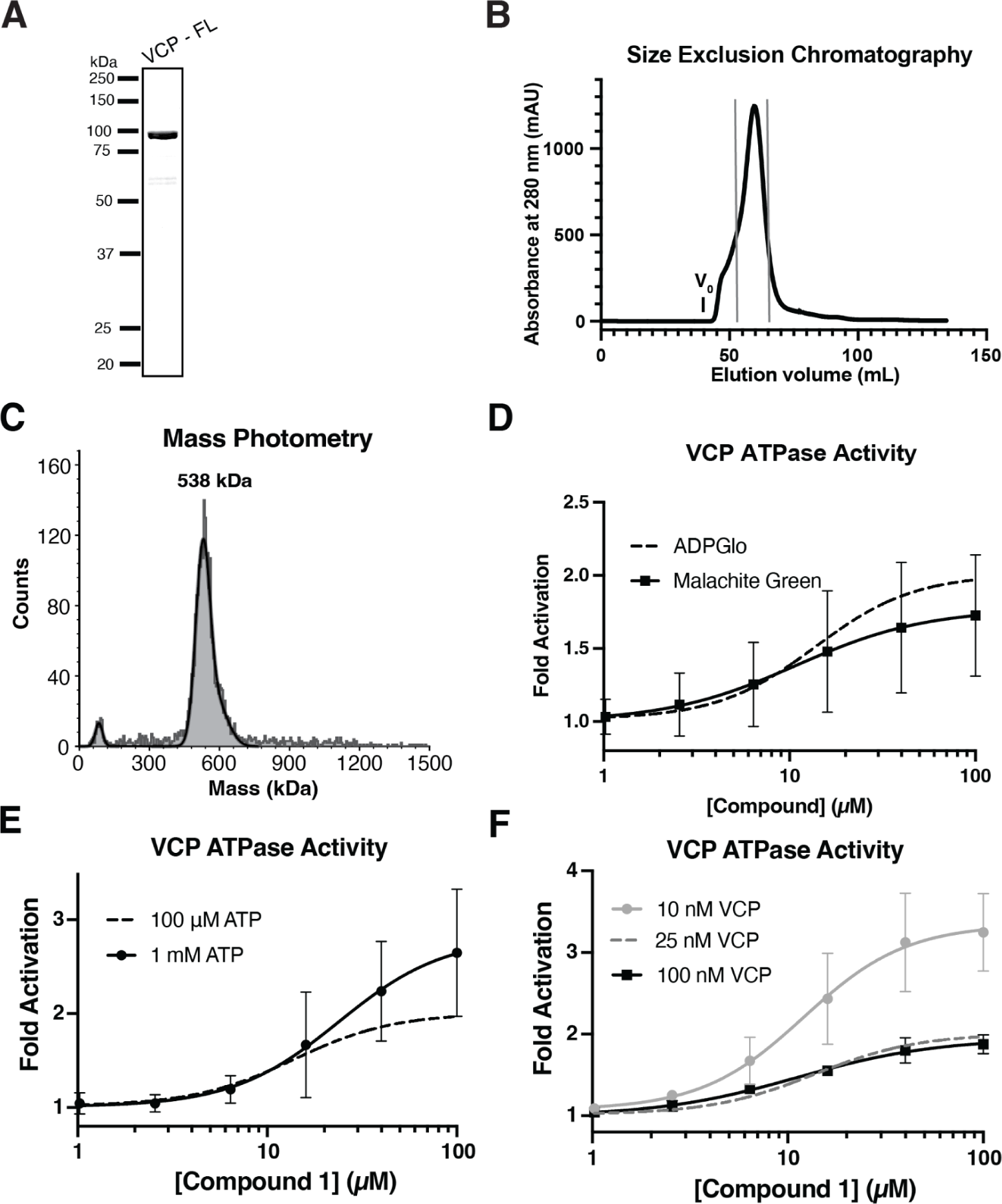
Characterizing VCP activity in the presence of Compound 1. (*A*) SDS-PAGE gel of purified recombinant VCP construct (Coomassie blue staining). (*B*) Absorbance at 280 nm for elution of recombinant VCP from a HiLoad 16/600 Superdex 200 column. Void volume is marked, and gray lines indicate collected sample for experiments. (*C*) Mass photometry for recombinant VCP, with mass for the largest peak indicated. A representative trace is shown (N = 3 independent experiments). (*D*) Concentration-dependent activation of the ATPase activity of VCP by Compound **1** (100 µM ATP, 1 hr endpoint colorimetric assay). Graph shows fold activation relative to DMSO control fit to a sigmoidal dose-response equation (mean +/− s.d., N = 3 independent experiments). Dotted line indicates fit from ADPGlo data reported in Fig. 1E. (*E*) Concentration-dependent activation of the ATPase activity of VCP by Compound **1** (1 mM ATP, 1 hr endpoint assay). Graph shows fold activation relative to DMSO control fit to a sigmoidal dose-response equation (mean +/− s.d., N = 3 independent experiments). Dotted line indicates fit from experiment performed at 100 µM ATP, reported in Fig. 1E. (*F*) Concentration-dependent activation of the ATPase activity of VCP by Compound **1** (100 µM ATP, 1 hr endpoint assay). Graph shows fold activation relative to DMSO control fit to a sigmoidal dose-response equation (mean +/− range, N = 2 independent experiments). Dotted line indicates fit from experiment performed at 25 nM VCP, reported in Fig. 1E.

**Fig. S2.**
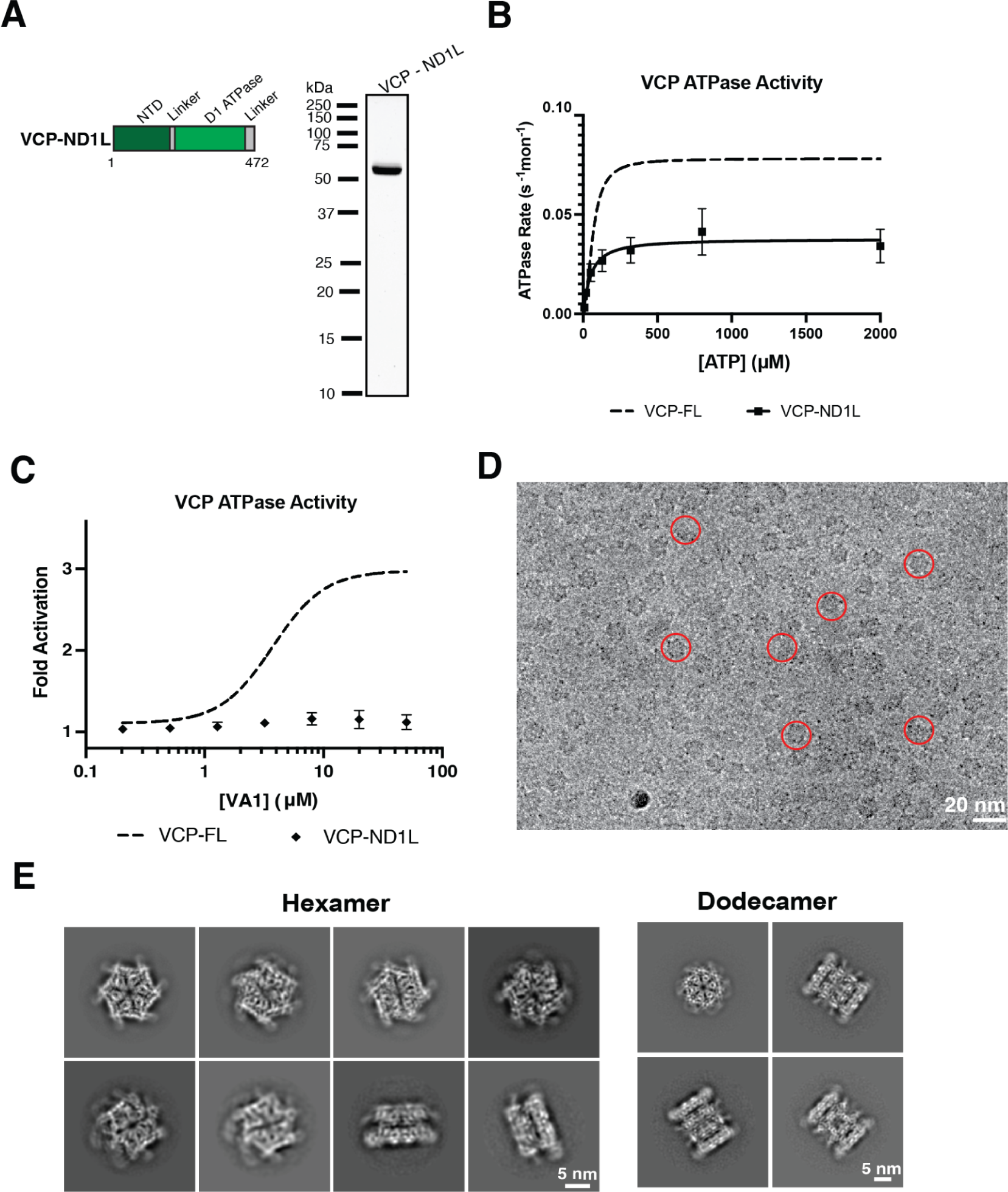
Structure-based analysis of the VCP/VA1 interaction. (*A*) Schematic showing the VCP-ND1L construct as a domain diagram (top, light gray box, not to scale, with the first and last residues and domains identified) (color-coded as in Fig. 1A) and SDS-PAGE gel of purified recombinant VCP-ND1L construct (Coomassie blue staining). (*B*) ATP concentration dependence of the steady-state activity of VCP-ND1L, analyzed using a Malachite Green assay. Rates were fit to the Michaelis-Menten equation for cooperative enzymes (mean +/− range, N = 2 independent experiments, dotted line indicates full-length VCP data from Fig. 1B). (*C*) Fold change in ATPase activity of VCP-ND1L in the presence of increasing concentrations of VA1 (100 µM ATP, 1 hr endpoint assay). Graph shows fold activation relative to DMSO control fit to a sigmoidal dose-response equation (mean +/− range, N = 2 independent experiments, dotted line indicates full-length VCP data from Fig. 2B). (D) Representative cryo-EM micrograph of VCP in the presence of VA1. Example particles of VCP are indicated (circles). (*E*) Representative 2D-class averages of VCP in the presence of VA1.

**Fig. S3.**
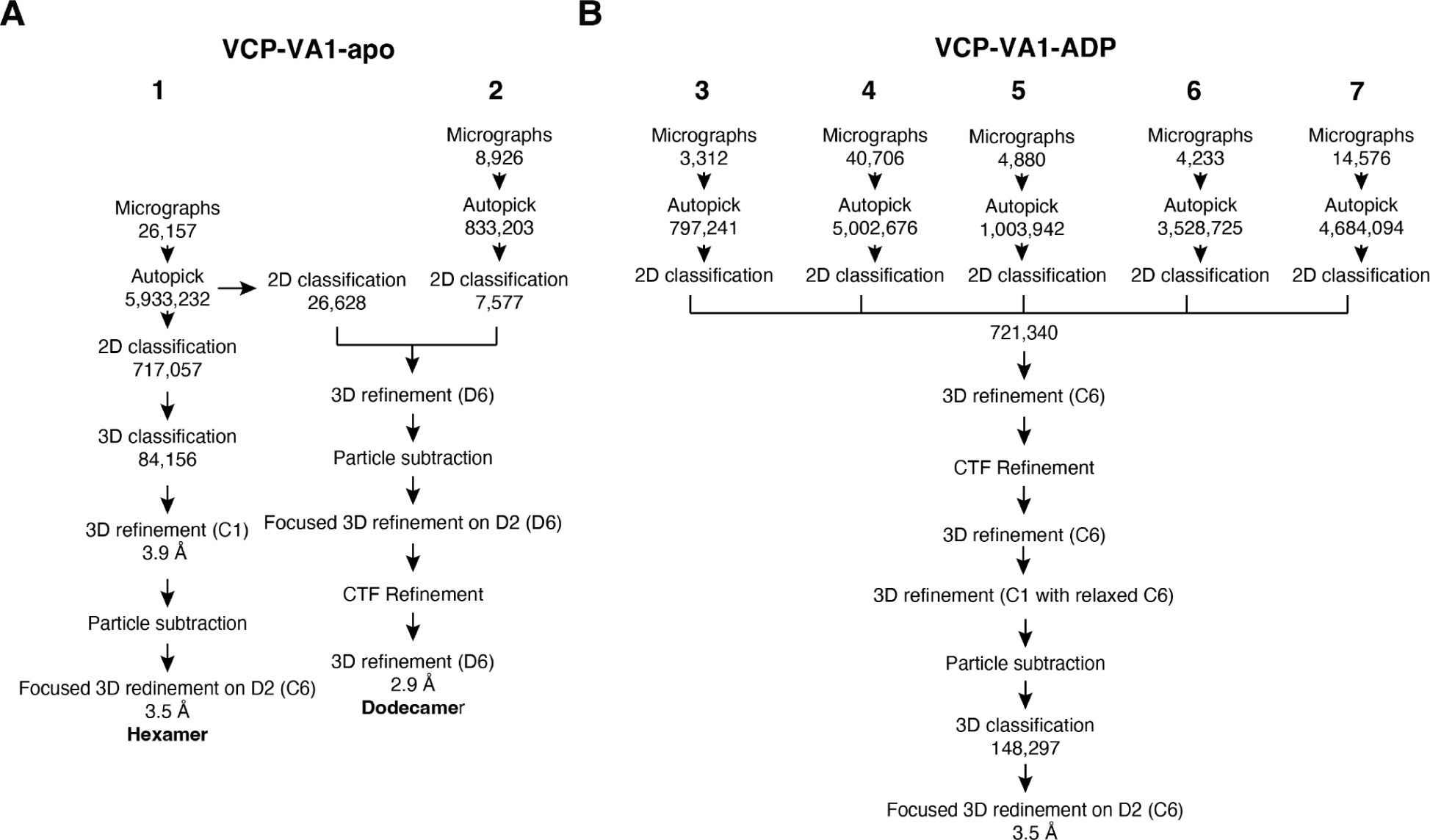
(*A-B*) Data processing workflow for reconstruction of VCP density maps and focused 3D classification and refinement procedures used to improve the resolution on the D2 domain density map, without addition of ATP (*A*) and with addition of ATP (*B*). Both conditions include the addition of VA1. Applied symmetry (D6 or C6) is indicated in parentheses.

**Fig. S4.**
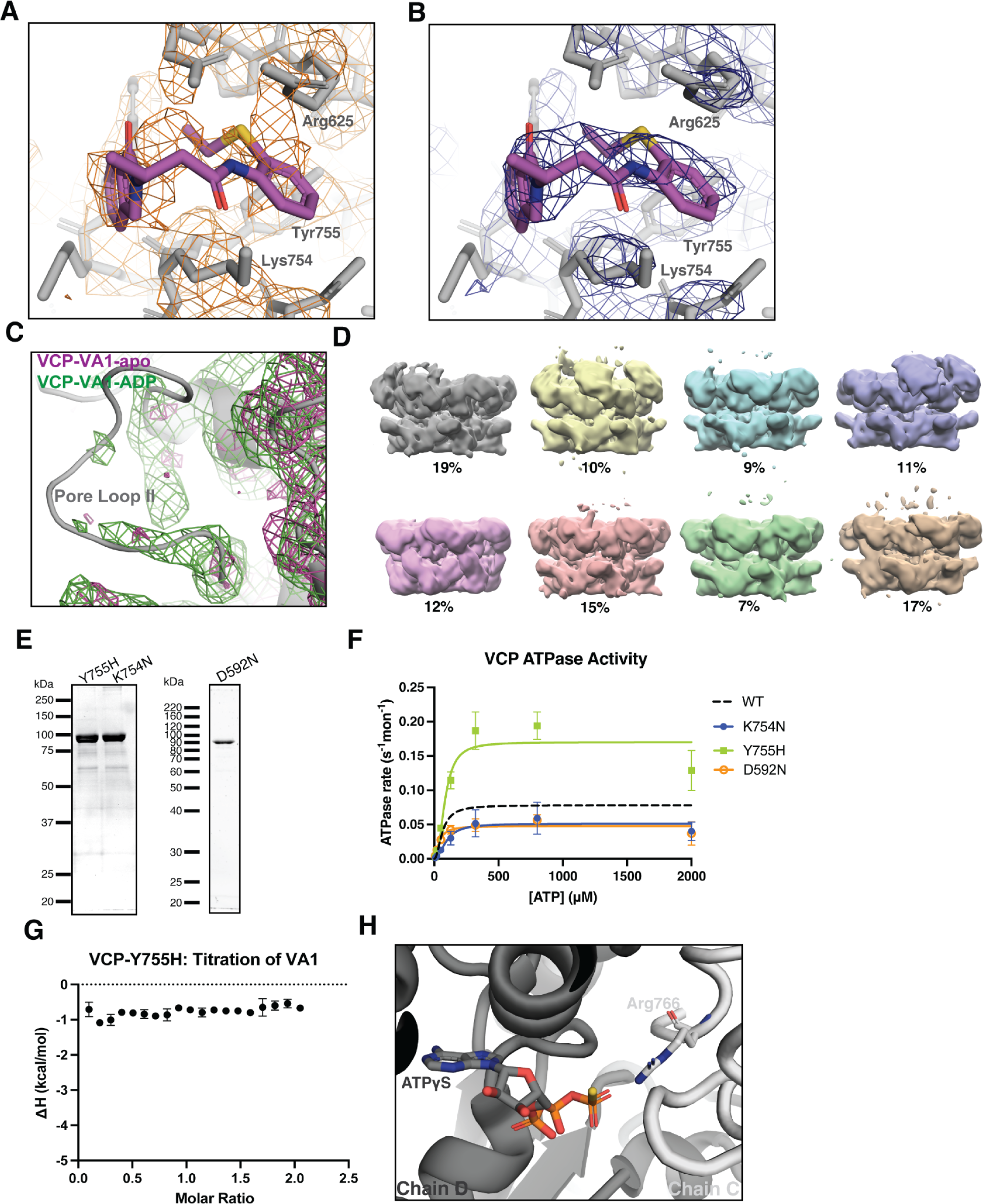
Examining the effects of nucleotide and point mutations on the VA1 binding site. (*A*) Model of VA1 in the proposed binding site (VA1, magenta, stick representation; VCP, gray, stick representation; VCP-VA1-apo density, orange, mesh). (*B*) Model of VA1 in the proposed binding site in the map of dodecamer (VA1, magenta, stick representation; VCP, gray, stick representation; density, blue, mesh). (*C*) View showing the D2 Pore Loop II, comparing maps for datasets with and without the addition of ATP (VCP, gray, stick representation; VCP-VA1-ADP map, green, mesh; VCP-VA1-apo map, magenta, mesh). (*D*) Representative 3D classes from VCP incubated with VA1 (50 µM) and ATP (100 µM). Particle distribution percentages are indicated below classes. (*E*) SDS-PAGE gels of purified recombinant VCP-K754N, -Y755H, and -D592N constructs (Coomassie blue staining). (*F*) ATP concentration dependence of the steady-state activity of VCP-K754N, -Y755H and -D592N mutants, analyzed using a Malachite Green assay. Rates were fit to the Michaelis-Menten equation for cooperative enzymes (mean +/− s.d., N = 3 independent experiments). Dotted line indicates fit from VCP-WT data shown in Fig. 1B. (*G*) Integrated data points from ITC-based analysis of VCP-Y755H in the presence of VA1 (mean +/− range, N = 2 independent experiments). (*H*) Model of VCP-ATPγS (PDB: 5FTN) (cartoon representation, Chain C in light gray, and Chain D in dark gray; ATPγS and Arg-766 shown in stick representation). View focuses on the D2 ATPγS binding site in Chain D.

**Table S1.**
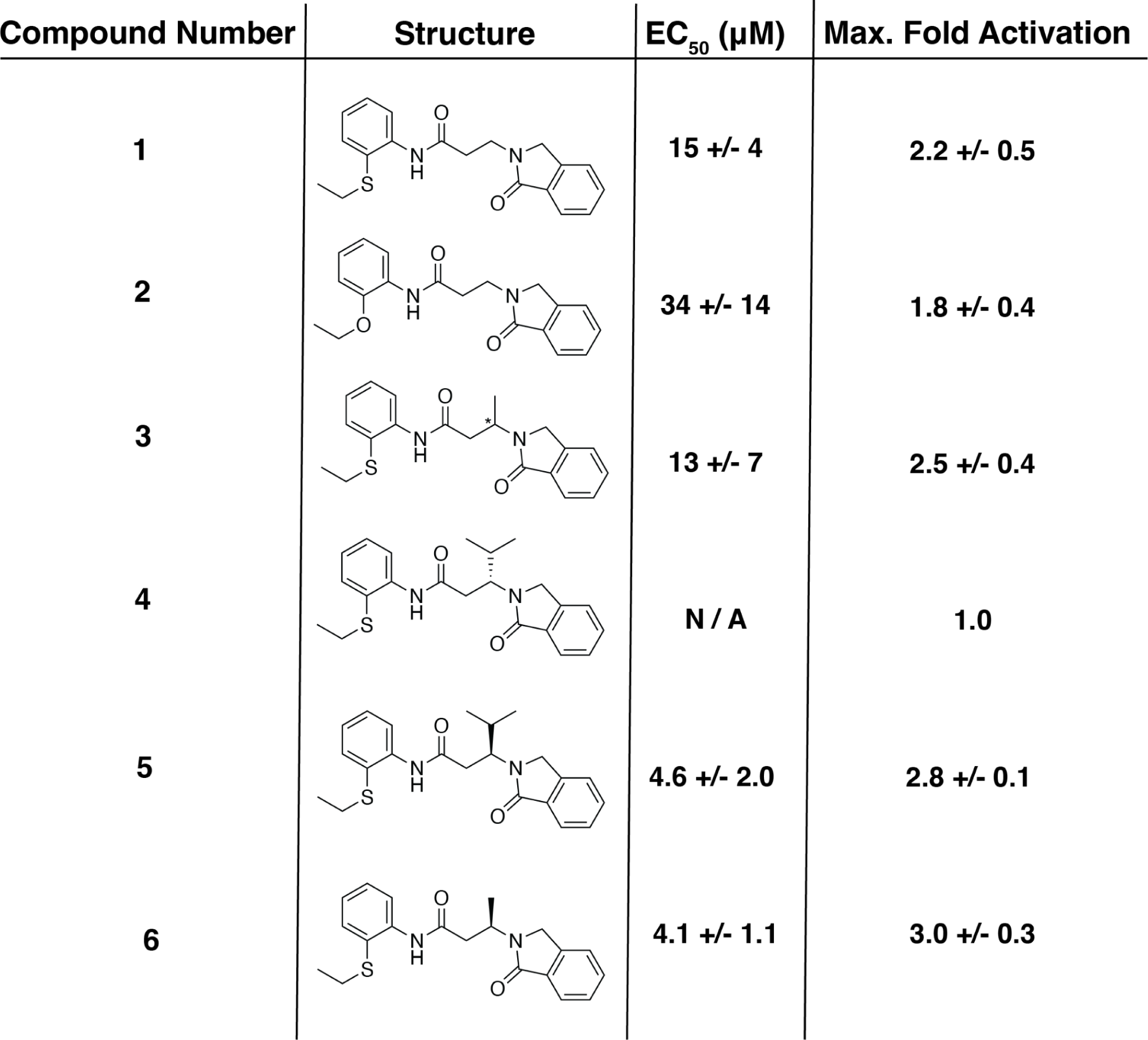
Analysis of VCP ATPase activity in the presence of Compound 1 analogs. Values for the maximum fold activation (saturated at high concentrations), as well as the EC_50_ of activation, are shown, analyzed using ADPGlo ATPase assays (100 µM ATP, 1 hr endpoint assay) (mean +/− s.d., N = 9 for Compound **1**, N = 7 for Compound **2**, N = 4 for Compounds **3** and **4**, N = 3 for Compound **5**, N = 5 for Compound **6**/VA1) (all replicates represent independent experiments).

| Compound Number | Structure | EC<sub>50</sub> (μM) | Max. Fold Activation |
| --- | --- | --- | --- |
| 1 |  | 15 +/- 4 | 2.2 +/- 0.5 |
| 2 |  | 34 +/- 14 | 1.8 +/- 0.4 |
| 3 |  | 13 +/- 7 | 2.5 +/- 0.4 |
| 4 |  | N / A | 1.0 |
| 5 |  | 4.6 +/- 2.0 | 2.8 +/- 0.1 |
| 6 |  | 4.1 +/- 1.1 | 3.0 +/- 0.3 |

**Table S2.** Cryo-EM data collection settings. Related to Fig. S3 and Materials and Methods.

|  | VCP-VA1-apo |  | VCP-VA1-ADP |  |  |  |  |
| --- | --- | --- | --- | --- | --- | --- | --- |
|  | 1 | 2 | 3 | 4 | 5 | 6 | 7 |
| Acceleration Voltage (kV) | 300 | 300 | 300 | 300 | 300 | 300 | 300 |
| Camera | K3 | K2 | K2 | K3 | K3 | K3 | K3 |
| CS Corrector | Yes | No | No | Yes | Yes | Yes | No |
| Energy Filter | Yes<br>(20 e- slit) | No | No | Yes<br>(20 e- slit) | Yes<br>(20 e- slit) | Yes<br>(20 e- slit) | Yes<br>(20 e- slit) |
| Exposure Time (s) | 1.25 | 6 | 5.2 | 1.3 | 2.5 | 2.5 | 2.5 |
| Frame Number | 25 | 20 | 13 | 20 | 30 | 30 | 30 |
| Pixel Size (Å/pixel) | 0.515 | 0.81 | 1.335 | 0.86 | 1.076 | 1.076 | 1.083 |
| Dose (electrons/Å<sup>2</sup>) | 49 | 27.4 | 23.5 | 20 | 51 | 51 | 48 |

